# Genomic diversity in felids correlates with range and density, not census size

**DOI:** 10.1101/2025.02.20.639334

**Authors:** Michaël P. Meeus, Jonas Lescroart, Hannes Svardal

## Abstract

As the world is hit by the sixth mass extinction, it becomes increasingly important to understand the factors relevant to the conservation of species, so that we may protect biodiversity to the best of our abilities. Although genetic diversity is known to reflect population demography and contribute to genetic health and adaptability, it is not explicitly used as a criterion in assessments by the International Union for the Conservation of Nature (IUCN). Additionally, studies comparing diversity estimates between species often rely on summarizing results across studies, which use different methodologies and may not be suited for direct comparison. Here we performed a family-wide assessment of genomic diversity in Felidae, covering most extant species. We tested for correlations between autosomal heterozygosity and ecological traits across (sub)species, and whether a subspecies’ genetic diversity was associated with its IUCN threat category. We found evidence for genetic diversity to be strongly positively correlated with both geographic range size and population density, but not with census size. Furthermore, although genetic diversity was not significantly correlated with IUCN status overall, threatened cat species had significantly lower levels of genetic diversity than non-threatened species. Our results confirm the association of population parameters and assessment of extinction risk with genetic diversity in one of the most iconic and threatened families of land carnivores. While mechanisms and causality behind these associations will need to be the subject of further investigation, our study adds further credence to the importance of incorporating genomic information in risk assessment and conservation efforts.

## Introduction

### Genetic diversity measures in service of conservation

Many species across the globe suffer reductions in population size and available habitat as a result of anthropogenic activity (Andermann et al. 2020). As these populations decline, so too does the genetic variation that exists within them (Willoughby et al. 2015). Genetic diversity is the key resource that allows species to adapt to novel conditions through natural selection. Loss of this genetic diversity will thus render species more vulnerable to extinction in the long term (Otto 2018). Declining populations are also likely to experience an increase in inbreeding, further reducing diversity in the gene pool, resulting in decreased fitness and reduced response to selection (Amos and Balmford 2001; Willi et al. 2022).

It is thus clear that genetic diversity is an important factor in the conservation of species and populations, and, accordingly, there have been increasing calls to collect more genetic data and increase the weight of genetic diversity in conservation assessments, something that is currently lacking in IUCN Red List assessments (Laikre et al. 2020; Hoban et al. 2020, 2024; Garner et al. 2020; Theissinger et al. 2023). Crucial topics of active debate are the relationship and potential predictive power of autosomal heterozygosity, a common measure of genetic diversity inferred from genomic data, with respect to a species’ threat status. Genomic data have been proven helpful in predicting a species’ threat status or conservation priority (Fernandez-Fournier et al. 2021; Wilder et al. 2023), but not universally (Teixeira and Huber 2021; Kuderna et al. 2023). The latter studies found that no clear relationship could be determined between heterozygosity and a species’ threat status across mammals and primates, respectively. However, it is equally possible that genetic data may be able to reveal species or populations at conservation risk that are currently not recognised using IUCN criteria (Willoughby et al. 2015).

### Application of genetic diversity measures in felid conservation

The cat family (Felidae) consists of at least 41 extant species, divided into two subfamilies, Felinae and Pantherinae (Kitchener et al. 2017) (Fig. S1). The family consists exclusively of hypercarnivores that fulfil important ecological roles in their environments, usually as mesopredators, while the larger species are generally apex predators and keystone species (Tensen 2018a, b). These top-down control effects, particularly from the larger apex predators, can exert substantial influences down the entire trophic chain (Ripple and Beschta 2006, 2008; Jorge et al. 2013; Sarasola et al. 2016). Beyond that, felids are typically charismatic species and score highly as favoured species for conservation with the public, making them suitable flagship species (Macdonald et al. 2015). Because of their ecological importance and charismatic appeal, felid research was an early adopter of using genomic data to further conservation research, as exemplified by studies on the Iberian lynx, Florida panther and cheetahs (Pimm et al. 2006; Johnson et al. 2010; Abascal et al. 2016; O’Brien et al. 2017; Theissinger et al. 2023).

Today, there is an increasing number of studies using autosomal heterozygosity to investigate members of the Felidae family, often in a conservation context, including the above mentioned Iberian lynx (Abascal et al. 2016), pumas (Saremi et al. 2019) and cheetahs (Prost et al. 2022), as well as jaguars (Lorenzana et al. 2022), lions (Armstrong et al. 2020; de Manuel et al. 2020), leopards (Paijmans et al. 2021; Pečnerová et al. 2021), species of the genus *Leopardus* (Lescroart et al. 2023), bobcats (Lin et al. 2022), snow leopards (Solari et al. 2023), tigers (Cho et al. 2013; Liu et al. 2018; Armstrong et al. 2021), black-footed cats (Yuan et al. 2024), and clouded leopards (Yuan et al. 2023).

### Study-specific design limits comparability of genetic diversity estimates across studies

A few metastudies have also looked at trends in genetic diversity within the broader family, rather than individual species or genera, using microsatellite data (Azizan and Paradis 2021; Azizan et al. 2023). These metastudies found that there is a relationship between genetic diversity and both population density (Azizan et al. 2023) and threat status of a species (Azizan and Paradis 2021). However, these studies have two limitations. First, traditional markers, such as microsatellites used in these studies, focus on a small, biased set of polymorphic loci that are not representative of the average genetic diversity in the genome (Väli et al. 2008). In comparison, whole-genome sequencing (WGS) data yields estimates of population genetic parameters with greatly increased accuracy, because it derives from the genome in its entirety (Allendorf 2017; Fuentes-Pardo and Ruzzante 2017). Availability of WGS data has been rapidly increasing over the past decade, and for cats specifically, this type of data is now publicly accessible for almost all species. Second, metastudies aggregate genetic diversity estimates from multiple individual studies. However, such a comparison may not be justified, if different methodologies are applied in different studies. This issue is especially apparent for estimates of heterozygosity from WGS data, which are sensitive not only to the choice of filter parameters and software, but also to the choice of reference genome (Brandt et al. 2015; Gopalakrishnan et al. 2017; Ros-Freixedes et al. 2018; Armstrong et al. 2020; Duchen and Salamin 2021; Prasad et al. 2022). Therefore, comparing estimates across studies may result in inaccurate conclusions and, instead, a single methodology should be applied to all genomic data. Some large-scale studies have carried out such analyses to look at whole-genome genetic diversity and how it relates to extinction risks in primates (Kuderna et al. 2023) and across placental mammals (Wilder et al. 2023). Previous studies applying such a unified approach to Felidae (Armstrong et al. 2020; Barnett et al. 2020; Lescroart et al. 2023; Solari et al. 2023) were limited in their taxonomic scope or the number of species included. As such, no study has applied a singular methodology to estimate and compare the heterozygosity across the majority of extant members of Felidae.

In the current study, we used previously published, raw whole-genome sequencing data to obtain estimates of heterozygosity for most members of the Felidae family (*sensu* Kitchener et al., 2017), covering all eight lineages of extant Felidae, excluding the domestic cat (*Felis catus*) and missing, notably, the Sunda leopard cat (*Prionailurus javanensis*) and African golden cat (*Caracal aurata*) (Fig. S1). A bay cat sample (*Catopuma badia*) was included in the initial dataset, but excluded from the final set of heterozygosity estimates due to poor data quality. While past studies were often focused on specific subgroups of felids, such as the genera *Panthera* (Armstrong et al. 2020), *Leopardus* (Lescroart et al. 2023) or “big cat species” (Solari et al. 2023), and some used a limited amount of species (∼15) across a wider taxonomic net (Barnett et al. 2020), our use of nearly all non-domesticated species of cats and a single methodological pipeline will allow, for the first time, for relatively unbiased comparisons between species across the Felidae family. We also compiled data on various species traits and population parameters at the species and subspecies level in order to uncover traits that may affect genetic diversity. We expected that larger population densities, geographic ranges and census sizes correlate positively with genetic diversity, as these parameters should relate to increased population mutation rate, reduced genetic drift and thus larger effective population sizes, an indicator of neutral genetic diversity (Frankham 2012; Leffler et al. 2012; Hague and Routman 2016; Myhre et al. 2016; Grundler et al. 2019; Jeon et al. 2024). On the other hand, we expected species with a larger body size or mass to have lower levels of genetic diversity than small-bodied species, because larger species typically maintain smaller population sizes (Silva et al. 2001; White et al. 2007; Grundler et al. 2019). Larger species are also more likely to suffer greater reductions in gene flow due to habitat fragmentation than do smaller species (Figueiredo et al. 2015). Finally, we compared genetic diversity to IUCN threat classification at subspecies level, testing the hypothesis that threatened subspecies have lower levels of genetic diversity.

### Materials and Methods

### Data collection

To enable comparison of genetic diversity across a broad range of felid species, we compiled a dataset consisting of 100 samples across 39 felid species. Samples were selected by querying the Sequence Read Archive and European Nucleotide Archive databases for short-read whole-genome sequencing data. We limited our query to samples of non-domesticated, extant Felidae species sequenced to medium or high depth of genomic coverage from fresh material (Table S1). Sequences and metadata were downloaded with fastq-dl v2.0.4 (Petit et al. 2023). Samples were classified at the subspecies level according to taxonomic and geographic information in the metadata. For the purpose of our analyses, any samples without sufficient metadata were assigned to the species’ most common subspecies; their uncertain origin is indicated where relevant. Classification followed the reference work *Felids and hyenas of the world* (José R. Castelló 2020), which itself largely follows the revised Felidae taxonomy published by the IUCN Cat Specialist Group (Kitchener et al. 2017), but distinguishes some additional populations of particular interest below the subspecies level. Samples were also labelled as captive, when obtained from, e.g., a zoo population, or non-captive when obtained from the wild (Table S2).

Data on species traits and population parameters (body mass, body size, gestation, litter size, longevity, census size, range and density) was compiled at the species and subspecies level by consulting the following sources: *Felids and hyenas of the world* (José R. Castelló 2020), the *Handbook of the mammals of the world* (Wilson and Mittermeier 2009), the IUCN Red List of Threatened Species (IUCN 2023) (Table S3), and information made available by the IUCN/SSC Cat Specialist Group (www.catsg.org). Body sizes refer to measurements from the tip of the muzzle to the base of the tail. Geographic ranges were extracted from spatial distribution data made available by the IUCN Red List of Threatened Species, excluding all polygons in the categories *Extinct* and *Possibly Extinct*. Subspecies ranges were obtained by subdividing the species ranges in QGIS v3.0 according to the geographic information listed in Kitchener et al. (2017). Surface areas were then calculated with custom Python code (see ‘Data Availability’).

### Mapping and variant calling

We aligned the raw reads of each sample to two reference genomes, applied filtering steps and called variant sites to obtain one set of genome-wide variants for each reference genome, using a custom pipeline (see ‘Data Availability’). We used two publicly available, highly contiguous genome assemblies as references, representing both subfamilies in Felidae: for Felinae, a domestic cat (*Felis catus*) genome (GenBank accession: GCA_018350175.1) (Bredemeyer et al. 2023) and for Pantherinae, a tiger (*Panthera tigris*) genome (GenBank accession: GCA_018350195.2) (Bredemeyer et al. 2023). This strategy allowed us to detect reference bias that may be introduced due to different genetic distances between the various samples and the reference genomes of both subfamilies (Prasad et al. 2022).

We created indices of the reference genomes using BWA v0.7.17 (Li and Durbin 2009) and SAMtools v1.14 (Danecek et al. 2021). Raw sequence reads of the sample set were then mapped to both reference genomes using the BWA-MEM algorithm. Using SAMtools, the aligned reads were tagged with mate scores and sorted by leftmost coordinates. We then marked duplicate reads, created an index for the Binary Alignment Map (BAM) file of each sample and obtained basic statistics for each alignment. BAM files were then compressed with Crumble v0.8.3 (Bonfield et al. 2019). We called bases jointly for all samples using BCFtools v1.14 (Danecek et al. 2021) and indexed the resulting Binary Call Format (BCF) file. We filtered the call set using BCFtools and custom Python code, retaining sites with mean mapping quality >50, <10% mapped reads with quality zero, allelic balance above 20%, depth of coverage <150% of the genome-wide average, and <20% missing genotype calls. We recorded the coordinates of all the sites that passed the filtering criteria using BEDTools v2.30.0 (Quinlan and Hall 2010) and further refer to this fraction of the genome as the ‘accessible genome’. We then selected all biallelic single nucleotide polymorphism (SNP) variants located in the accessible genome from the call set, which were stored in Variant Call Format (VCF).

### Heterozygosity Estimates

For each sample in the VCF file, we counted the number of homozygous, heterozygous and missing genotypes in each autosome with a custom Python script (see ‘Data Availability’). We calculated heterozygosity as the number of heterozygous sites, taking into account a corresponding fraction of missing genotypes, and normalizing our count with the size of the accessible genome. Variance around the mean heterozygosity of each sample was calculated as the weighted standard deviation of estimates obtained from each of the 18 autosomes. Specifically, our estimator of heterozygosity is given as:

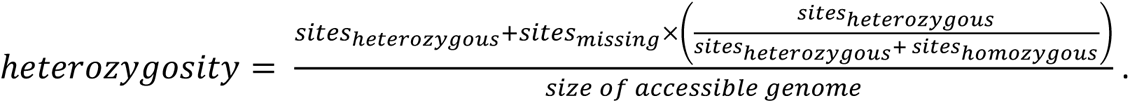

### Statistical analysis

We performed statistical analyses to test for biases in our heterozygosity estimates and to detect correlations between heterozygosity and various species and population characteristics or threat status. All analyses were conducted using custom R code (see ‘Data Availability’). Exploration of the data revealed a significant correlation between mean coverage and heterozygosity best explained by a power correlation of the formula: *y* = 0.006251 ∗ *x*^−0.556245^ (adjusted R^2^=0.343; p <0.001), with the samples with the lowest coverage showing high heterozygosity (Fig. S2). When compared to our overall heterozygosity estimates and to other samples of the same species, the estimates for these low-coverage samples appeared inflated. We therefore excluded samples with an average coverage below 7 for both reference genomes from further analysis (Cat_bad_PBA2; Lyn_ruf_mLynRuf1.p; Her_yag_HJA5, Aci_jub_EH_ID75, Aci_jub_DZ_ID42 and Aci_jub_AJU173) (Table S4; see Table S5 for a full list of samples and datapoints removed from various parts of the analysis).

We explored potential effects of reference bias in our heterozygosity estimates by calculating the strength of correlation between the estimates derived from mapping to the two distinct reference assemblies. In addition, we directly compared the estimate of each sample obtained from mapping to its phylogenetically closest reference (same subfamily) to the estimate obtained from the more distant reference (different subfamily).

To enable regression analysis, heterozygosity was averaged at the subspecies level for body mass, body size and geographical range, while for gestation time, litter size and lifespan, estimates were averaged at species level. For population density and census size, data was averaged at the subspecies level where possible, and at the species level otherwise. We then performed a simple linear regression to test the correlation between heterozygosity, the dependent variable, and each ecological variable separately, as the independent variable. P-value, sample size (n), correlation coefficient (R) and the coefficient of determination (adjusted R^2^) of the linear regression were added to the plots using ggpmisc v0.6.1 (Aphalo 2024). Lastly, we produced diagnostic plots for each correlation to verify adherence to the model assumptions, particularly looking for signs of heteroscedasticity and potentially influential outliers, as these will have the greatest effect on model accuracy (Schmidt and Finan 2018). Outliers were considered influential when the Cook’s distance exceeded 5 (Table S5). We redid the linear regression after transforming the data or removing influential outliers, where appropriate.

We also performed a multiple regression analysis in order to assess the relative predictive value of each ecological variable in relation to heterozygosity. First, we averaged the heterozygosity estimates of the individual samples by subspecies and computed simple linear regression models for each ecological variable. Again, where appropriate we performed a transformation of the data or removed outliers based on diagnostic plots. Next, we assessed intercorrelation between the ecological variables using the Kendall rank correlation coefficient (Kendall’s Tau), visualised with the GGally package v2.2.1 (Schloerke et al. 2024). We then constructed a multiple regression model with the variables that showed significant correlation with heterozygosity, but not with any other variable included in the model. The lack of multicollinearity of independent variables in the model was verified by calculating the variance inflation factor using the car package v3.1-3 (Fox and Weisberg 2019). Lastly, we determined the relative predictive value of each ecological variable in the multiple regression model by calculating their partial coefficients of determination using the Sensemakr v0.1.6 package (Cinelli et al. 2024).

To evaluate the difference in heterozygosity between threat categories, we used the IUCN Red List Categories classification and averaged heterozygosity by subspecies before applying a Welch’s t-test to compare between threatened (Vulnerable, Endangered and Critically Endangered) and non-threatened (Least Concern and Near Threatened). We also applied an ANOVA test followed by Tukey’s Honest Significant Difference test to evaluate differences between the separate IUCN categories. Finally, we explored potential differences between samples obtained from captive or wild-collected individuals by comparing the two groups across the entire family and within the *Panthera* lineage using Welch’s t-tests. The latter lineage was chosen for its large representation of both groups compared to the other lineages. Results were visualised with boxplots. Plots were generated using the ggplot2 (v3.5.1) and ggpubr (v0.6.0) packages (Wickham 2016; Kassambara 2023). The significance threshold for all analyses was set at p < 0.05.

## Results

### Mapping and base calling

We mapped the raw sequencing reads of each sample (94 million - 1 billion reads per sample, see Table S1) to both reference genomes and applied a series of filter steps, resulting in accessible genome sizes of 78% (1.78 Gbp) of the domestic cat assembly and 80% (1.82 Gbp) of the tiger assembly. Base calling yielded270 and 275 million variant sites in the total sample set after filtering. The number of variants retained per sample varied between 1.9-35.2 million biallelic SNPs (Table S6).

### Minor effect of reference genome on heterozygosity estimates

Heterozygosity estimates obtained with reference assemblies of both subfamilies were highly correlated (R = 1.00; p < 0.001) (Fig. 1a). Estimates were also consistent when comparing between those using a phylogenetically close and more distantly related reference genome (Fig. 1b). In those cases where estimates were noticeably different, it was typically an increase in heterozygosity when mapped to the distant reference genome (Fig. 1c). This occurred primarily in tigers (*Panthera tigris*) and members of the wildcat lineage (genus *Felis*), particularly the African wildcat (*F. lybica*) (Fig. S3). The largest difference was found in an Asiatic wildcat individual (FLI4) and constitutes an absolute increase in heterozygosity by 0.047 percentage points, or a 21.7% increase when mapped to the more distant reference genome relative to the closer reference (Table S7). Overall, our results suggest that with our variant filtering the choice of reference genome is of minor importance at the phylogenetic scale of this study. Because of the high similarity between both sets of heterozygosity estimates, the remainder of the analysis was performed using the domestic cat set.

**Fig. 1.**
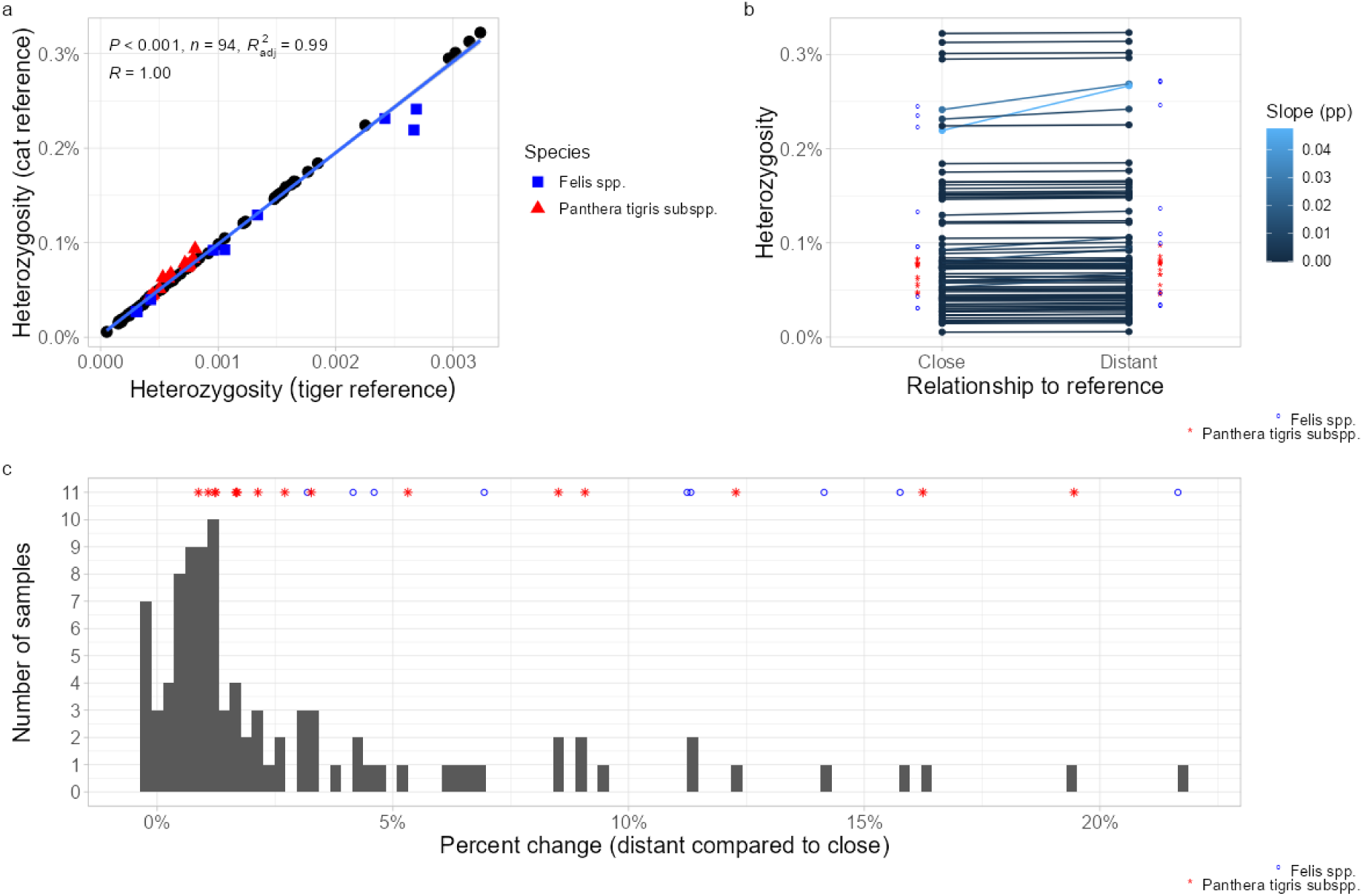
Effect of using different reference genomes on heterozygosity estimates. **a** Correlation between heterozygosity estimates using a reference genome belonging to a tiger and a domestic cat. **b** Estimates of heterozygosity obtained from mapping against the reference genome from the same subfamily versus the reference genome from the other subfamily. Data points derived from the same sample are connected with a line, the slope of which indicates percentage point difference. **c** Histogram showing the difference in heterozygosity estimates given as a percent increase from estimates derived from the close reference to estimates derived from the distant reference

### 54-fold variation in heterozygosity estimates across the cat family

Our estimates of heterozygosity range from 0.006% (SD = 0.002%) in a sample from the Indian population of lions, *Panthera leo leo,* to 0.322% (SD = 0.012%) in an ocelot, *Leopardus pardalis*, revealing a 53.7-fold difference across the cat family (Fig. 2, grouped by subspecies, Fig. S4, grouped by species). Other species with high levels of heterozygosity are the serval (*Leptailurus serval*) (0.295%), and African/Asiatic wildcats (*Felis lybica lybica/ornata*) (0.219-0.231%), while other species with notably low levels of heterozygosity were the Andean cat (*Leopardus jacobita*) (0.014%) and the snow leopard (*Panthera uncia*) (0.017 - 0.019%).

**Fig. 2.**
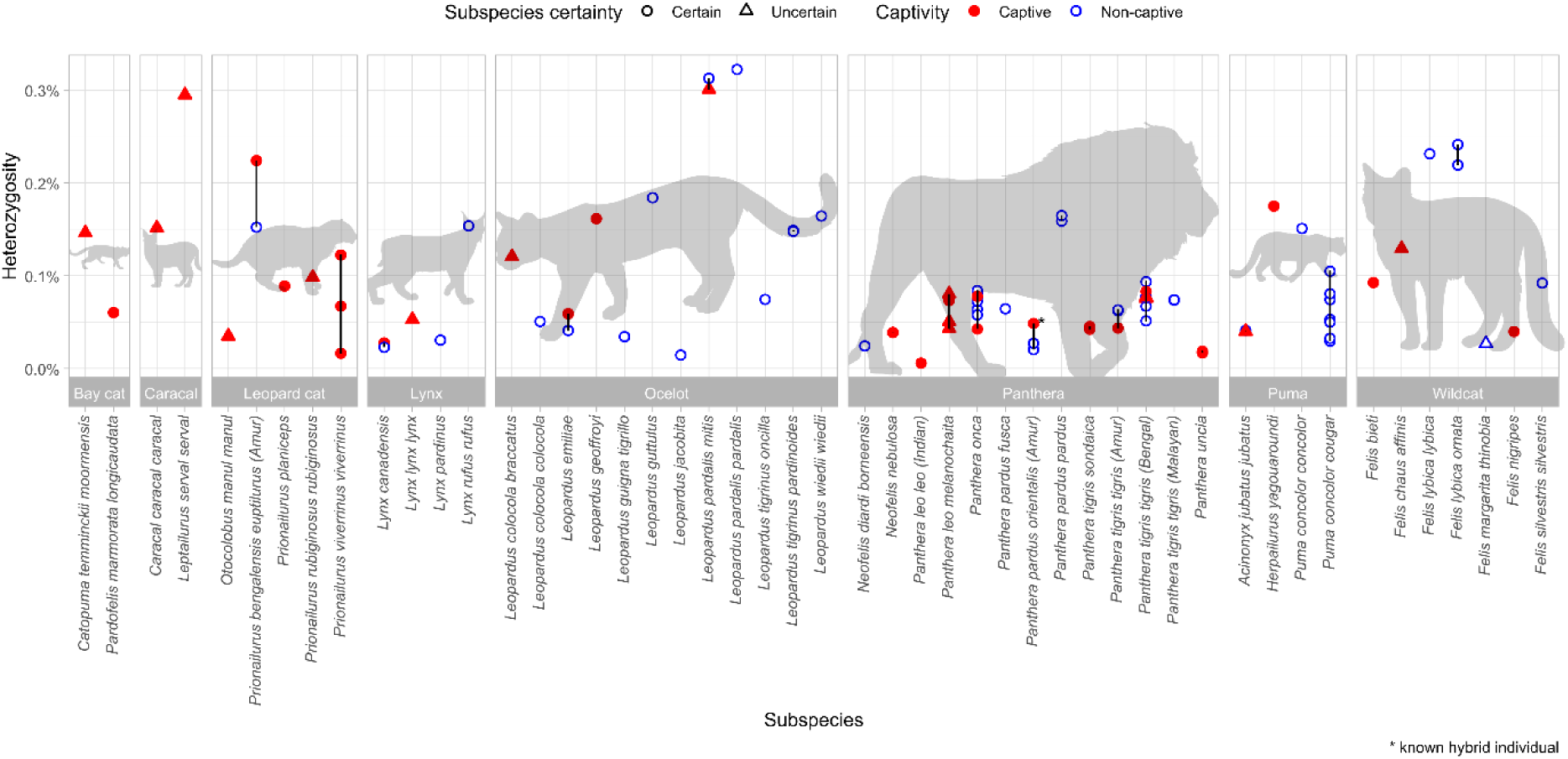
Estimates of heterozygosity for all samples using the cat reference genome, grouped per lineage and subspecies. Red filled shapes denote samples from captive populations; blue open shapes denote samples from wild populations. Circles denote samples where subspecies classification is certain; triangles denote samples where subspecies classification is uncertain. Silhouettes obtained from PhyloPic (www.phylopic.org)

We found no significant difference in heterozygosity estimates between captive and non-captive samples across the entire family (t_(89.4)_ = -0.927; p = 0.357), but within the Panthera lineage, there was a significant difference (t_(39.2)_ = -2.83; p = 0.007), with captive samples exhibiting lower genetic diversity on average (Fig. S5). Heterozygosity values vary greatly both across and within the different felid lineages, and vary noticeably even between samples of the same species. For instance, heterozygosity differs between African leopards and the Asiatic subspecies (resp., *Panthera pardus pardus* and *P. p. orientalis/fusca*), between North/Central and South American pumas (resp., *Puma concolor cougar* and *P. c. concolor*), and within North/Central American pumas (*P. c. cougar*) (Fig. 2). Of the eight suprageneric lineages commonly referred to in Felidae, none was uniformly high or low in genetic diversity.

### Heterozygosity is significantly correlated with geographic range size and population density

The simple linear regressions (Fig. 3) indicated a clear positive relationship between heterozygosity and range (R = 0.47; p < 0.001). Population density initially showed no significant correlation with heterozygosity (R = 0.15; p = 0.369). Outlier analysis, however, revealed the presence of two strong outliers, the guigna (*Leopardus guigna*) and jungle cat (*Felis chaus*) (Fig. S6 a-d). Removal of the outliers revealed a strong positive correlation with heterozygosity (R = 0.52; p = 0.001) (Table S5). Alternatively, using a more conservative estimate for guigna population density also resulted in a significant positive relationship (R = 0.36; p = 0.025) (Fig. S7).

**Fig. 3.**
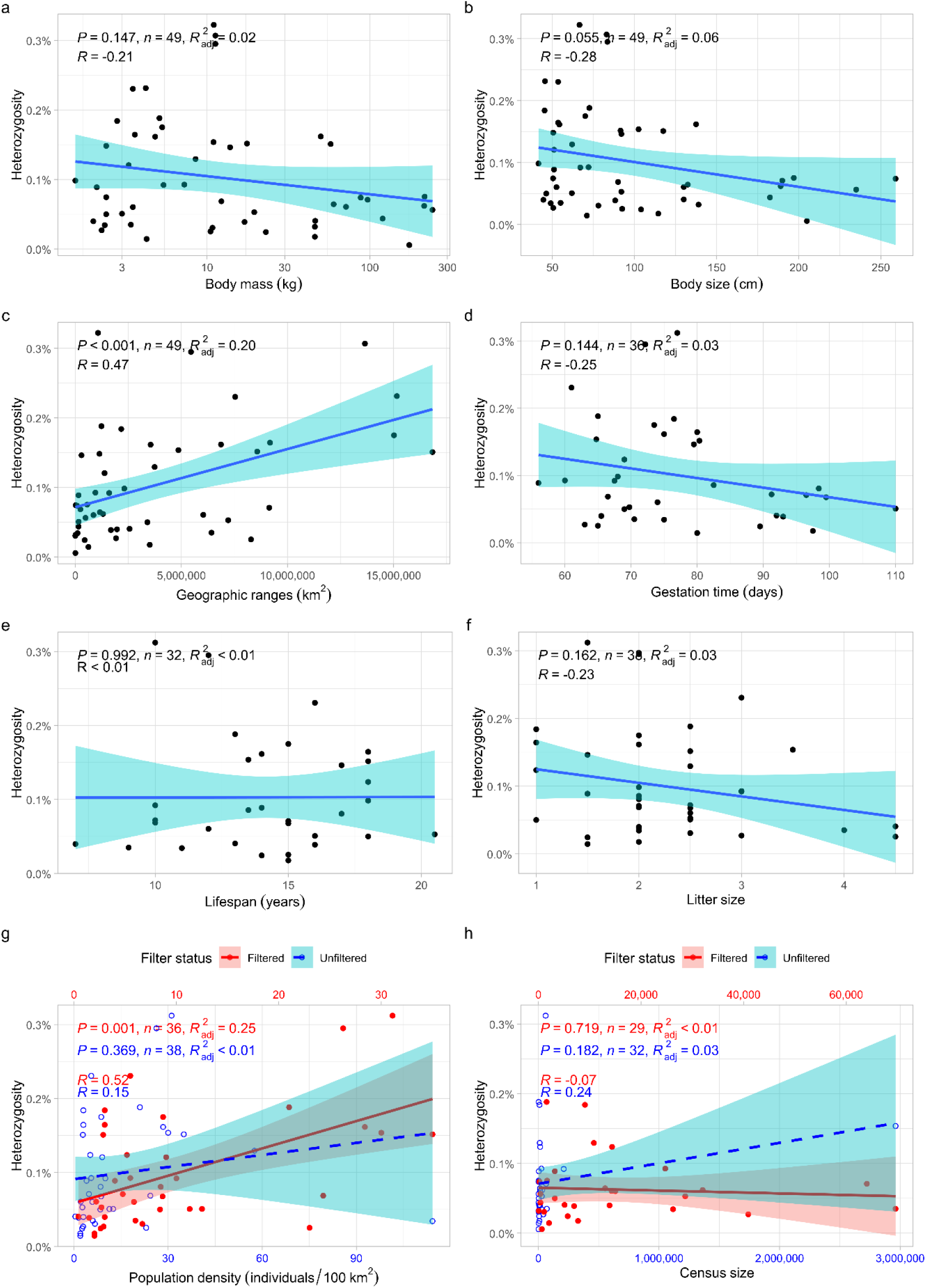
Simple linear regressions between heterozygosity and ecological traits. Heterozygosity as a function of **a** log-transformed body mass, **b** body size, **c** geographic range, **d** average gestation time, **e** average lifespan, **f** average litter size, **g** average population density, **h** average census size. Averaging is at subspecies level (**a-c**), species level (**d-f**), or both depending on data availability (**g-h**). **g** and **h** show regression with (blue; open circles; dashed line) and without (red; filled circles; solid line) outliers. Datapoints were transformed (red) and imposed on a secondary axis (red; top) for clarity

Census size showed no significant relationship with heterozygosity, both before (R = 0.24; p = 0.182) and after outlier removal (R = -0.07; p = 0.719) (Fig. 3h; see also Fig. S6 e-h and Table S5). Average lifespan, also, showed no correlation with heterozygosity (R < 0.01; p = 0.99). Litter size (R = -0.23; p = 0.162), body size (R = -0.28; p = 0.055), body mass (log-transformed) (R = -0.21; p = 0.147) and gestation time (R = -0.25; p = 0.144) all showed weak, non-significant negative relationships with heterozygosity (Fig. 3).

To build the multiple regression model, we selected only the independent variables that significantly correlated with heterozygosity at the subspecies level and lacked significant correlation with other such independent variables (Fig. S8). This resulted in a model with two explanatory variables: range and population density (τ_B_ range∼density = 0.040 (filtered; p = 0.98); 0.003 (conservative; p = 0.73)). Because the population density data was less complete at the subspecies level than the geographic range data, we summarized the range data to match the sample size of the density data (see: “Statistical analysis”). The model explains 29.3% of the variation in heterozygosity (adjusted R^2^ = 0.2933). The low Variance Inflation Factor (VIF = 1.001) confirmed absence of multicollinearity in the model. Range was a more important predictor (partial R^2^ = 0.230; p = 0.003) compared to population density (partial R^2^ = 0.155; p = 0.016) when the conservative density estimate was used for guigna. However, density increased in importance when the guigna datapoint was omitted entirely (Range: partial R^2^ = 0.201; p = 0.007, Density: partial R^2^ = 0.275, p = 0.001). The overall model that omitted the guigna and jungle cat datapoints also explained more of the variation in heterozygosity (adjusted R^2^ = 0.3845; p < 0.001; VIF = 1.017).

### Heterozygosity is significantly lower in threatened species

There was a noticeable trend of declining heterozygosity estimates going from Least Concern to Critically Endangered categories (with the exception of Critically Endangered showing slightly higher heterozygosity than Endangered) (mean heterozygosity: LC = 0.162%; NT = 0.099%; VU = 0.075%; EN = 0.042%; CR = 0.050%). Overall, heterozygosity estimates differed significantly between IUCN categories (F_(4;44)_ = 5.91; p < 0.001). Pairwise comparisons revealed that significant differences only existed between subspecies classified as Least Concern and Vulnerable (p = 0.006) and Least Concern and Endangered (p = 0.002) (Fig. 4a). When splitting the data into threatened and non-threatened categories, heterozygosity was higher on average in the non-threatened (0.144%) than in the threatened (0.063%) subspecies (p < 0.001) (Fig. 4b).

**Fig. 4.**
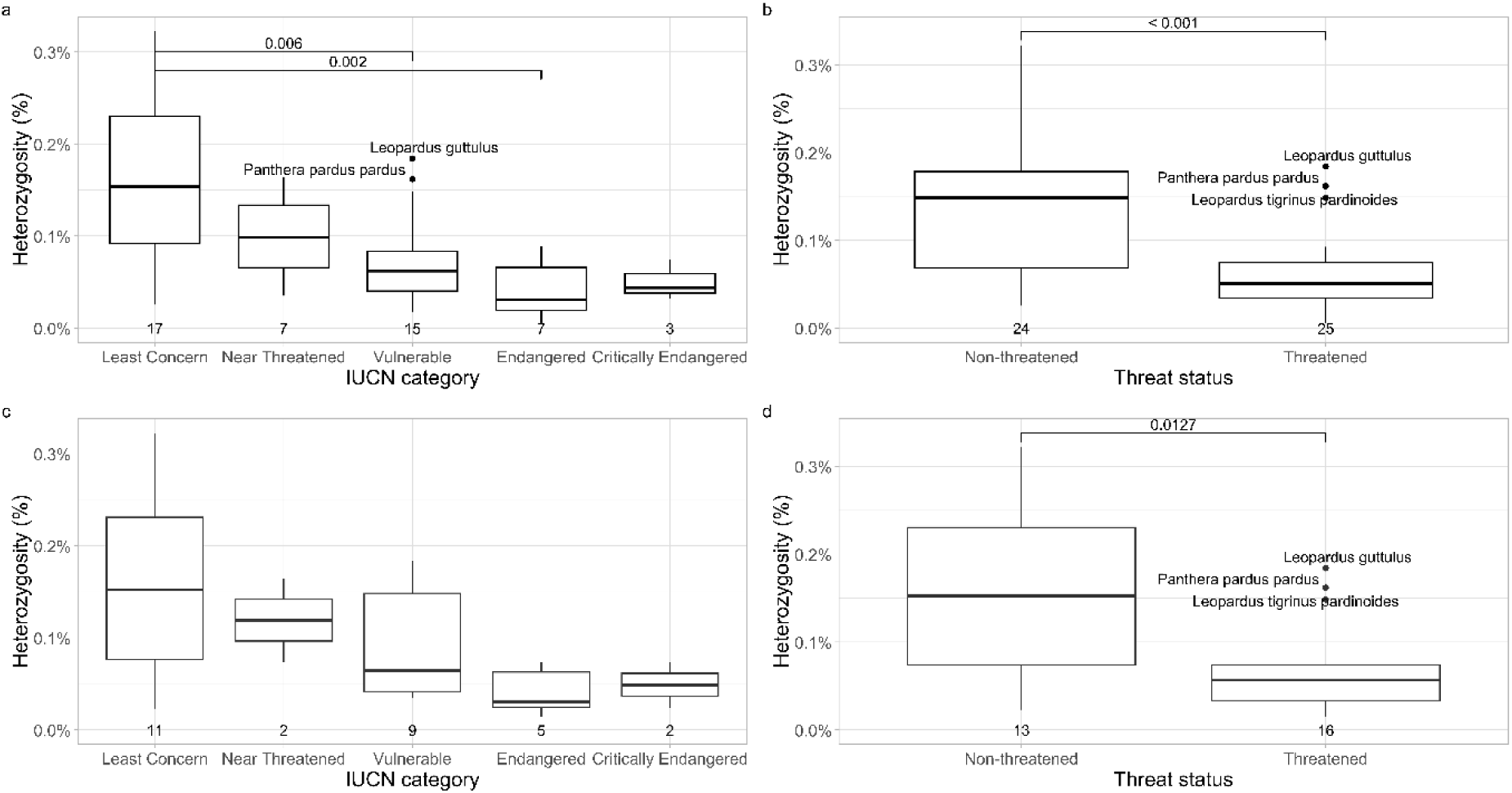
Differences in average heterozygosity of subspecies between **a** the different IUCN categories, **b** threatened and non-threatened subspecies, **c** the different IUCN categories using only non-captive samples, **d** threatened and non-threatened subspecies using only non-captive individuals. Comparisons between threat categories were carried out using a Welch’s t-test. Comparisons between IUCN categories were carried out using a Tukey HSD test after rejection of an ANOVA test. Sample sizes per category are shown below each box

Heterozygosity did not exhibit significant differences between IUCN categories when only non-captive individuals were considered (p = 0.066). The difference between threatened and non-threatened subspecies, however, remained significant (p = 0.0127) (Fig. 4c & 4d).

## Discussion

### Choice of reference genome has little effect on heterozygosity estimates in felids

Reference bias may affect estimates of genetic diversity in complex and contrasting ways, dependent on features of the genome assembly and the phylogenetic relation of the reference species to the focal taxa. Increased phylogenetic distance to the reference genome may both inflate (Prasad et al. 2022) or underestimate (Degner et al. 2009; Sousa and Hey 2013; Ros-Freixedes et al. 2018; Duchen and Salamin 2021) heterozygosity through misalignments between the reference and the sample of interest. Estimates may further be impacted by the quality of the reference genome (Gopalakrishnan et al. 2017; Prasad et al. 2022) or the software and filtering strategy employed (Brandt et al. 2015; Duchen and Salamin 2021).

Considering all these caveats, it becomes clear that comparisons of estimates obtained from different studies may very well lead to inaccurate results. To address these issues, we subjected WGS data of nearly all extant Felidae species to a single heterozygosity estimation pipeline, thereby ensuring identical treatment of each sample in terms of mapping, quality filtering and estimation of heterozygosity. Our resulting set of heterozygosity estimates (Fig 2, Table S4; see tables S7 & S8 for different layouts) offer a largely unbiased view of the relative differences in genetic diversity found in individual samples across the cat family. Furthermore, our use of two high-quality, ultra-contiguous assemblies, representing both subfamilies, (*Felis catus* with N50 of 148.5 Mbp and 39 gaps, and *Panthera tigris* with N50 of 146.9 Mbp and 65 gap), allowed to assess potential reference bias, both in terms of quality and phylogenetic distance to the samples of interest.

We found that estimates were generally consistent between reference genomes (Fig. 1), contrasting previous studies (Armstrong et al. 2020; Prasad et al. 2022). This may be the result of the high quality of both reference genomes and the high level of synteny among felid genomes (Davis et al. 2009; Cho et al. 2013; Armstrong et al. 2020). Clear signs of reference bias, considered as >5% difference between same-sample estimates (n=18; see Table S7), mostly concerned samples closely related to one of the reference species (n=12), i.e., tigers (*Panthera tigris*) which share a common ancestor 7.5-9.2 kya (Armstrong et al. 2021) and species in the domestic cat lineage (*Felis spp.*) with a common ancestor 3.34 mya and ubiquitous post-speciation gene flow (Yuan et al. 2024). Large relative differences between references may also be an artefact of low absolute heterozygosity (likely in samples of fishing cat and Canadian lynx; n=3), possibly compounding true reference bias (*Panthera* samples with low diversity, i.e., Asiatic lion and snow leopard; n=3). Where biased, heterozygosity was generally overestimated with the more distant reference, in line with the findings of Prasad et al. (2022), who surmised that this might be the result of misalignments.

### Genetic diversity across Felidae

The sample with the lowest heterozygosity (0.006 ± 0.002%) among our set of felids was an Asiatic lion (*Panthera leo leo*; formerly *P. l. persica*), consistent with previous findings (Azizan and Paradis 2021). The Asiatic lion once roamed across most of India, its range possibly extended as far west as Anatolia (Jhala et al. 2019). Habitat loss and hunting during the second half of the 19^th^ century eventually reduced the Asiatic lion population to less than 50 individuals in the Gir forest in India (Jhala et al. 2019). Its negative population trend has since been reversed as a result of conservation measures implemented during the 20^th^ century, elevating the species from Critically Endangered to Endangered status (Jhala et al. 2019). Regardless, the severe bottleneck the populations experienced is more than likely still showing its effects in the estimates of heterozygosity determined in this study. Other (sub)species with severely depleted levels of heterozygosity (<0.03%) were the Andean cat (*Leopardus jacobita*), snow leopard (*Panthera uncia*), Amur leopard (*Panthera pardus orientalis*), Canada lynx (*Lynx canadensis*), Bornean clouded leopard (*Neofelis diardi borneensis*) and Arabian sand cat (*Felis margarita thinobia*). These (sub)species have previously been reported to harbour exceedingly low genetic diversity (Paijmans et al. 2021; Bursell et al. 2022; Lescroart et al. 2023; Solari et al. 2023; Yuan et al. 2024), with the exception of Canada lynx. Studies on the Canada lynx have typically found high levels of heterozygosity (Campbell and Strobeck 2006), although heterozygosity was lower in peripheral and insular populations (Schwartz et al. 2003; Prentice et al. 2019). Another species from the Lynx lineage, the Iberian lynx (*Lynx pardinus*), is often brought forward as an example of extremely low heterozygosity among vertebrates (Abascal et al. 2016). With an estimate of 0.031 ± 0.005% for Iberian lynx, our family-wide comparison demonstrates that there are several other felids species that have lower levels of genetic diversity.

The highest estimates of heterozygosity in this study all belong to the ocelot (*Leopardus pardalis*), which is in concordance with past studies that have looked at genetic diversity in *Leopardus* and other felid species (Ramirez et al. 2022; Lescroart et al. 2023). Plausible explanations for the high diversity of the ocelot are its occurrence at high densities and broad distribution as well as higher levels of gene flow between populations compared to larger species such as jaguars (*Panthera onca*) (Figueiredo et al. 2015; Lescroart et al. 2023). Other (sub)species with high heterozygosity values (> 0.2%) include the serval (*Leptailurus serval*), the leopard cat (*Prionailurus bengalensis*) and the African/Asiatic wildcat (*Felis lybica lybica/ornata)*. The leopard cat and both the African and Asiatic wildcat have previously been shown to have high degrees of genetic diversity compared to other cat species (Barnett et al. 2020; Azizan and Paradis 2021; Yuan et al. 2024), while we were unable to find any earlier estimates of heterozygosity for the serval. It should also be noted that both the serval and the leopard cat sample were derived from captive individuals, which may affect genetic diversity.

The incorporation of multiple samples per species or, where available, per subspecies, revealed relatively uniform heterozygosity in some species and considerable intraspecific variation in others. Species such as ocelot, African wildcat, jaguar and tiger all exhibited low variation across samples, while high variation was observed in puma and leopard samples, in accordance with the findings of the studies that originally published the data (Kim et al. 2016; Saremi et al. 2019; Paijmans et al. 2021). Several other species showed considerable variation in heterozygosity, such as the fishing cat (*Prionailurus viverrinus*) (Fig. 2). However, as for some species we are partly or entirely dependent on data from captive individuals with little information on pedigree or origin, it is not possible to ascertain whether this variation is representative for natural populations, or the result of in- or outbreeding in captivity. For instance, the amur leopard (*Panthera pardus orientalis*) with the highest heterozygosity was confirmed to be a 30% hybrid with a North Chinese leopard (*P.p. orientalis, formerly P.p. japonensis*) (Kim et al. 2016). In other instances, such as the puma samples, the variation matches natural patterns of population structure. The highest level of heterozygosity in this species (0.15%), belongs to the South American puma (*Puma concolor concolor*), which has larger and better-connected populations than their northern conspecifics. In North America, populations were drastically reduced as a result of habitat destruction and hunting, until regulations were put in place during the mid-20^th^ century (Saremi et al. 2019). Genetic diversity in North American pumas ranges from 0.03% in an inbred and threatened population in Florida (Pimm et al. 2006; Johnson et al. 2010; Saremi et al. 2019), up to 0.08% and 0.10% in larger and outbred populations, respectively (Saremi et al. 2019). In general, diversity in puma populations, appears to match their degree of isolation (Riley et al. 2014; Saremi et al. 2019).

### Heterozygosity correlates with population density and geographic range

We tested for correlations between average heterozygosity of subspecies and various ecological traits of the subspecies using simple linear regression. We found that a subspecies’ heterozygosity was highly positively correlated with geographic range (Fig. 3c), conform with some studies that have previously found significant relationships between these two variables in *Drosophila* as well as threatened mammals (Leffler et al. 2012; Doyle et al. 2015), but in contrast with findings from Romiguier et al. (2014) and Azizan and Paradis (2021). Romiguier et al. examined a broad host of metazoan taxa, whereas the current study is limited to the Felidae family. Our observed correlation between heterozygosity and range may break down at larger phylogenetic scales owing to differences in life history traits or other aspects of biology that do not vary within a single family. On the other hand, Azizan and Paradis (2021) focused specifically on Felidae, yet did not observe a correlation between range and genetic diversity, which is likely a consequence of our different treatment of the spatial data available through the IUCN Red List. In their study, the authors used the range data as it is presented, i.e., at the level of species, whereas we carried out manual division of the spatial polygons to instead conform to the ranges of subspecies. Our approach allows for a more nuanced correlation analysis of geographic range and genetic diversity data, because the subspecies level more accurately approximates the population structure that shapes genetic diversity.

Average population density initially showed no significant correlation with genetic diversity values, in contrast with expectations based on earlier findings (Azizan et al. 2023). Exploring the data revealed the presence of two outliers (Fig. S6a-c), removal of which results in a significant positive relationship (Fig. 3g). These outlier data points belong to the guigna (*Leopardus guigna*) and jungle cat (*Felis chaus*), both having relatively low levels of genetic diversity and high estimates for their population density. The outliers are indicative of a general limitation of the population density data: a lack of estimates from different regions and/or populations. For both species, the number of available estimates is limited and both have estimates derived from pristine habitats (Belousova 1993; Dunstone et al. 2002), where one can expect density to be higher and thus not representative for the situation across the species’ range. Alternatively, using another, possibly more conservative estimate for the guigna also resulted in a significant positive relationship (Fig. S6d & S7), bringing it more in line with the findings of Azizan et al. (2023). Our finding of a positive relationship between population density and genetic diversity in felids aligns with the prediction from neutral theory and is in agreement with empirical observations in other taxa, e.g., in lizards (Hague and Routman 2016).

We also found that there were no significant relationships (P > 0.05) between genetic diversity and body size, or genetic diversity and body mass, unlike what was reported in earlier studies across metazoan taxa (Romiguier et al. 2014), butterflies (Mackintosh et al. 2019), mammals (Wooten and Smith 1985; Lino et al. 2019), tanagers (Brüniche-Olsen et al. 2019) and birds in general (Eo et al. 2011), but in agreement with an earlier study on felids (Azizan and Paradis 2021) and another in mammals (Doyle et al. 2015). It thus appears that measures of body size are not strong predictors of genetic diversity in felids. That said, we do observe a negative trend in felids between genetic diversity and body size (R = -0.28) or body mass (R = -0.27), as did Azizan and Paradis (2021). As argued by these authors, measures of body size are associated to other factors that might have causal links with heterozygosity. Indeed, larger-bodied species tend to have lower population sizes, a phenomenon known as Damuth’s rule (Damuth 1981), and thus generally lower N_e_, as well as lower rates of molecular evolution (Martin and Palumbi 1993; Eo et al. 2011). Interestingly, measures of body size did not show any notable correlation with the two significant variables – population density and geographic range (Fig. S8). This may be the result of relatively low sample sizes and lack of availability of population density data. On the other hand, relationships between measures of body size and both population density and geographic range have been found to be more complex than expected (Silva et al. 2001; White et al. 2007; Willig et al. 2009; Smith and Lyons 2013; Lyons et al. 2019).

Census size also showed no correlation with autosomal heterozygosity, contrary to our hypothesis that larger population sizes are linked to increased genetic diversity, as it has previously been shown that census size can be used as a proxy for effective population size with an N_e_/N ratio of around 0.11-0.14 (Frankham 1995, 2012; Palstra and Ruzzante 2008). Some past studies did manage to uncover a significant correlation with genetic diversity (Frankham 2012; Hague and Routman 2016), but many others were similarly unable to establish such correlation (Nabholz et al. 2008; Perry et al. 2012; Leffler et al. 2012). A possible explanation for the lack of correlation is the nature of the census data. The census data here represents the sum total of all mature individuals of a species or subspecies, which may be far greater than the true census sizes of individual, reproductively connected populations, thus masking potential correlations. Additionally, some studies have found evidence of a negative relationship between N_e_/N and N, particularly in very small populations, which may further muddle the relationship (Palstra and Ruzzante 2008; Myhre et al. 2016). A more fine-grained analysis with accurate estimations of census data at the population level would thus greatly help in uncovering how population size relates to genetic diversity in cats.

Finally, we found no relationship between genetic diversity and the life history traits of average litter size, lifespan and gestation time. Gestation time and litter size did both show a noticeable negative trend, however. The negative trend shown by gestation time is likely related to the negative trend noticed in body size and mass, as gestation time has been shown to be strongly correlated to body mass in carnivores (Danis and Rokas 2024). The negative trend in litter size on the other hand conflicts with earlier findings by Romiguier et al. (2014), who found a strong positive correlation between measures of genetic diversity and fecundity. It should be noted, however, that Romiguier et al. used a large variety of metazoan taxa, and thus had a much larger range of fecundity data compared to our taxonomically limited scale. Their study also revealed a negative correlation with lifespan where we found none. This too might be explained by the difference in taxonomic scope and the associated variability in lifespans.

A multiple linear regression model, which was created using the variables that were significant in the simple linear regression analysis (population density and geographic range), suggested that population geographic range was a better predictor of heterozygosity than population density, unlike the conclusion by Azizan et al. (2023), who identified population density as the main predictor of genetic diversity in felids. Factors leading to these different outcomes are likely our use of genome-wide estimates of diversity, our improved resolution in range data and inclusion of a higher number of species. It should be noted that the relative importance of both variables is reversed when the guigna data is removed. Future studies investigating these relationships would benefit greatly from improved estimates of population characteristics, especially population density.

### Threatened species have significantly lower levels of autosomal heterozygosity

We tested the relationship between genetic diversity and IUCN threat status within a single family, *Felidae*, using whole-genome sequence data. When deciding on conservation priorities, genome-wide variation is thought of as a suitable proxy for variation with adaptive potential, as demonstrated in yellow warblers and lodgepole pines (Fernandez-Fournier et al. 2021). Our results indicate that heterozygosity was not a good predictor of the different IUCN threat categories, although a negative trend was observed between heterozygosity and severity of the threat level, consistent with general expectations (Fig. 4a). These findings confirm those of earlier studies where heterozygosity was found to be a poor predictor of IUCN status (Teixeira and Huber 2021; Schmidt et al. 2023; Kuderna et al. 2023), while also supporting findings that there is a general, weak relationship between declining heterozygosity and increasing extinction risk (Schmidt et al. 2023). We also found that estimates of heterozygosity performed better as a predictor of extinction risk when the categories were summarized as “threatened” (LC, NT) and “non-threatened” (VU, EN, CR), as was the case in previous studies (Willoughby et al. 2015; Azizan and Paradis 2021; Wilder et al. 2023; Kuderna et al. 2023).

There are several possible reasons for the lack of congruence between the genetic data and IUCN classification. Most importantly, it takes time for genetic diversity to reflect changes in a population’s demography. Current estimates of heterozygosity may not capture recent, sudden population decline or fragmentation and instead represent historic population size and connectivity (Willoughby et al. 2015; Schmidt et al. 2023). The IUCN Red List also does not make use of genetic diversity data in its assessment criteria, meaning that species that might qualify for a threatened status from a genetic point of view are potentially overlooked by the IUCN criteria (Willoughby et al. 2015; Schmidt et al. 2023). Finally, the non-threatened categories, LC and NT, tend to have a much wider range of heterozygosity values compared to the threatened categories, VU, EN and CR, as well as more species listed, as has been reported elsewhere (Schmidt et al. 2023).

We have shown above that the high variability of genetic diversity estimates in North American pumas can be explained by the conditions and demographic history of the populations that each of the samples belong to. The wide variation in heterozygosity estimates present in non-threatened categories may therefore in part result from the sampling of threatened populations of an otherwise non-threatened (sub)species. For example, prior to the revision of the Felid taxonomy by Kitchener et al. (2017), the Florida puma, which exhibits the lowest levels of heterozygosity among puma samples here, was categorised as an endangered subspecies and currently still is listed as an endangered subpopulation (Nielsen et al. 2016).

### Genomic diversity in conservation

We have shown that in felids threatened species have significantly lower levels of genetic diversity than non-threatened species and that this heterozygosity can be linked to various species or population traits, among which are geographical range and population density, presumably because these traits reflect the effective population size to some degree (Myhre et al. 2016; Birzu et al. 2019; Jeon et al. 2024). In that regard, the lack of correlation between genetic diversity and census size in our sample set was unexpected, as lower population sizes are expected to lead to increased genetic stochasticity and inbreeding, and thus lower N_e_. However, as discussed above, this may simply be a limitation of the available data. The IUCN Red List criteria focus largely on census size, geographic distribution and the reduction thereof to establish whether a species belongs in a threatened category (IUCN Standards and Petitions Committee 2024).

Therefore, it appears the lack of correlation between census data and heterozygosity compromises the predictive capability of genetic diversity in classifying a species or subspecies as threatened, while any legitimate predictive power relies at least partly on geographic distribution as a common determinant of both genetic variation and threat assessment.

Our results suggest that genetic diversity may serve as a useful diagnostic tool in future conservation assessments, with the caveat that changes in genetic diversity are delayed compared to the immediate changes in population parameters such as geographic range or population size (Epps and Keyghobadi 2015; Gargiulo et al. 2024). Beyond its utility as a measure for monitoring population viability, genetic diversity is critically important in its own right to maintain healthy populations capable of adapting to new threats (Slate et al. 2000; Reed and Frankham 2003; Charpentier et al. 2005; Markert et al. 2010; Takahashi et al. 2018; Clarke et al. 2024). Low levels of genetic diversity have repeatedly been linked to various health issues in wild cat populations. In pumas, genetic impoverishment has been shown to result in kinked tails, cryptorchidism and teratospermia (Huffmeyer et al. 2022). Similarly, cheetahs were also found to have malformed sperm alongside an overall lower sperm count, as well as a lack of variation in the immune system (O’Brien et al. 2017). These examples illustrate the relevance of measuring genetic diversity for wildlife conservation, and our heterozygosity estimates may help to identify the felid species and subspecies that are most at risk.

Overall, our study bolsters the call for genomic data and diversity measures to constitute an essential aspect of conservation efforts. To improve future analyses, several issues warrant attention, particularly regarding data availability and bias. For many of the samples used in this study, metadata detailing the sample’s origin was hard to find or unavailable, making it difficult to correctly assign subspecies labels and associated species characteristics, limiting the accuracy of the analyses. Furthermore, there exists a clear bias in data availability (genetic and non-genetic) and studies towards the larger and often more recognisable species in the family (Brodie 2009; Pérez-Irineo and Santos-Moreno 2013; Zanin et al. 2015; Tensen 2018a; Azizan and Paradis 2021). Additional field and genomic research on the small and inconspicuous species of cats will allow for a better understanding of the processes that determine genetic diversity and extinction risk in cats and by, extension, other taxa, so that we may push for increasingly competent prioritization strategies to better preserve global biodiversity.

## Acknowledgments

The authors gratefully acknowledge support of the Research Fund of the University of Antwerp. M.P.M. was supported by a University Research Fund (BOF – Seal of Excellence) awarded by the University of Antwerp. J.L. was supported by Research Foundation — Flanders (grant number 1128621N). Special thanks to Alexandros Bantounas for providing code to compute the surface area of spatial polygons. We thank staff at the high-performance computing core facility CalcUA for assistance in using computing infrastructure and are grateful for the computational resources made available by the Flemish Supercomputer Centre. Unpublished genome assemblies and sequencing data for a black-footed cat, an African lion, a jaguar, a snow leopard, a leopard cat and a fishing cat are used with permission from the DNA Zoo Consortium (dnazoo.org).

## Statements & Declarations

### Author Contributions

CRediT author statement — Michaël Patrick Meeus: Data Curation, Formal Analysis, Methodology, Software, Writing - original draft. Jonas Lescroart: Conceptualization, Methodology, Validation, Writing - review and editing. Hannes Svardal: Supervision, Writing - review and editing.

### Data Availability

We accessed and re-used publicly available whole-genome sequencing data (Table S1). Custom R and Python code used in this study are available on an archived GitHub repository [pending: https://doi.org/10.5281/zenodo.14597830]. Variant call sets (VFC format) and spatial polygons of subspecies ranges (shape files) are shared on Figshare [pending: link to data sets].

## Reference List

Abascal F, Corvelo A, Cruz F, et al (2016) Extreme genomic erosion after recurrent demographic bottlenecks in the highly endangered Iberian lynx. Genome Biology 17:251. 10.1186/s13059-016-1090-1

Allendorf FW (2017) Genetics and the conservation of natural populations: allozymes to genomes. Molecular Ecology 26:420–430. 10.1111/mec.13948

Amos W, Balmford A (2001) When does conservation genetics matter? Heredity 87:257–265. 10.1046/j.1365-2540.2001.00940.x

Andermann T, Faurby S, Turvey ST, et al (2020) The past and future human impact on mammalian diversity. Science Advances 6:eabb2313. 10.1126/sciadv.abb2313

Aphalo PJ (2024) ggpmisc: Miscellaneous Extensions to “ggplot2”

Armstrong EE, Khan A, Taylor RW, et al (2021) Recent Evolutionary History of Tigers Highlights Contrasting Roles of Genetic Drift and Selection. Molecular Biology and Evolution 38:2366– 2379. 10.1093/molbev/msab032

Armstrong EE, Taylor RW, Miller DE, et al (2020) Long live the king: chromosome-level assembly of the lion (Panthera leo) using linked-read, Hi-C, and long-read data. BMC Biology 18:3. 10.1186/s12915-019-0734-5

Azizan A, Anile S, Nielsen CK, et al (2023) Population density and genetic diversity are positively correlated in wild felids globally. Global Ecology and Biogeography n/a: 10.1111/geb.13727

Azizan A, Paradis E (2021) Patterns and drivers of genetic diversity among Felidae species. Biodivers Conserv 30:519–546. 10.1007/s10531-020-02103-5

Barnett R, Westbury MV, Sandoval-Velasco M, et al (2020) Genomic Adaptations and Evolutionary History of the Extinct Scimitar-Toothed Cat, Homotherium latidens. Current Biology 30:5018–5025.e5. 10.1016/j.cub.2020.09.051

Belousova AV (1993) Small Felidae of Europe Asia and Far East. Lutreola 2:16–21

Birzu G, Matin S, Hallatschek O, Korolev KS (2019) Genetic drift in range expansions is very sensitive to density dependence in dispersal and growth. Ecology Letters 22:1817–1827. 10.1111/ele.13364

Bonfield JK, McCarthy SA, Durbin R (2019) Crumble: reference free lossy compression of sequence quality values. Bioinformatics 35:337–339. 10.1093/bioinformatics/bty608

Brandt DYC, Aguiar VRC, Bitarello BD, et al (2015) Mapping Bias Overestimates Reference Allele Frequencies at the HLA Genes in the 1000 Genomes Project Phase I Data. G3 Genes|Genomes|Genetics 5:931–941. 10.1534/g3.114.015784

Bredemeyer KR, Hillier L, Harris AJ, et al (2023) Single-haplotype comparative genomics provides insights into lineage-specific structural variation during cat evolution. Nat Genet 55:1953– 1963. 10.1038/s41588-023-01548-y

Brodie JF (2009) Is research effort allocated efficiently for conservation? Felidae as a global case study. Biodivers Conserv 18:2927–2939. 10.1007/s10531-009-9617-3

Brüniche-Olsen A, Kellner KF, DeWoody JA (2019) Island area, body size and demographic history shape genomic diversity in Darwin’s finches and related tanagers. Molecular Ecology 28:4914–4925. 10.1111/mec.15266

Bursell MG, Dikow RB, Figueiró HV, et al (2022) Whole genome analysis of clouded leopard species reveals an ancient divergence and distinct demographic histories. iScience 25:105647. 10.1016/j.isci.2022.105647

Campbell V, Strobeck C (2006) Fine-scale genetic structure and dispersal in Canada lynx (Lynx canadensis) within Alberta, Canada. Can J Zool 84:1112–1119. 10.1139/z06-099

Charpentier M, Setchell JM, Prugnolle F, et al (2005) Genetic diversity and reproductive success in mandrills (Mandrillus sphinx). Proceedings of the National Academy of Sciences 102:16723– 16728. 10.1073/pnas.0507205102

Cho YS, Hu L, Hou H, et al (2013) The tiger genome and comparative analysis with lion and snow leopard genomes. Nat Commun 4:2433. 10.1038/ncomms3433

Cinelli C, Hazlett C, Ferwerda J (2024) sensemakr: Sensitivity Analysis Tools for Regression Models

Clarke JG, Smith AC, Cullingham CI (2024) Genetic rescue often leads to higher fitness as a result of increased heterozygosity across animal taxa. Molecular Ecology 33:e17532. 10.1111/mec.17532

Damuth J (1981) Population density and body size in mammals. Nature 290:699–700. 10.1038/290699a0

Danecek P, Bonfield JK, Liddle J, et al (2021) Twelve years of SAMtools and BCFtools. GigaScience 10:giab008. 10.1093/gigascience/giab008

Danis T, Rokas A (2024) The evolution of gestation length in eutherian mammals. Proceedings of the Royal Society B: Biological Sciences 291:20241412. 10.1098/rspb.2024.1412

Davis BW, Raudsepp T, Pearks Wilkerson AJ, et al (2009) A high-resolution cat radiation hybrid and integrated FISH mapping resource for phylogenomic studies across Felidae. Genomics 93:299–304. 10.1016/j.ygeno.2008.09.010

de Manuel M, Barnett R, Sandoval-Velasco M, et al (2020) The evolutionary history of extinct and living lions. Proceedings of the National Academy of Sciences 117:10927–10934. 10.1073/pnas.1919423117

Degner JF, Marioni JC, Pai AA, et al (2009) Effect of read-mapping biases on detecting allele-specific expression from RNA-sequencing data. Bioinformatics 25:3207–3212. 10.1093/bioinformatics/btp579

Doyle JM, Hacking CC, Willoughby JR, et al (2015) Mammalian genetic diversity as a function of habitat, body size, trophic class, and conservation status. Journal of Mammalogy 96:564–572. 10.1093/jmammal/gyv061

Duchen P, Salamin N (2021) A Cautionary Note on the Use of Genotype Callers in Phylogenomics. Systematic Biology 70:844–854. 10.1093/sysbio/syaa081

Dunstone N, Durbin L, Wyllie I, et al (2002) Spatial organization, ranging behaviour and habitat use of the kodkod (Oncifelis guigna) in southern Chile. Journal of Zoology 257:1–11. 10.1017/S0952836902000602

Eo SH, Doyle JM, DeWoody JA (2011) Genetic diversity in birds is associated with body mass and habitat type. Journal of Zoology 283:220–226. 10.1111/j.1469-7998.2010.00773.x

Epps CW, Keyghobadi N (2015) Landscape genetics in a changing world: disentangling historical and contemporary influences and inferring change. Molecular Ecology 24:6021–6040. 10.1111/mec.13454

Fernandez-Fournier P, Lewthwaite JMM, Mooers AØ (2021) Do We Need to Identify Adaptive Genetic Variation When Prioritizing Populations for Conservation? Conserv Genet 22:205–216. 10.1007/s10592-020-01327-w

Figueiredo MG, Cervini M, Rodrigues FP, et al (2015) Lack of Population Genetic Structuring in Ocelots (Leopardus pardalis) in a Fragmented Landscape. Diversity 7:295–306. 10.3390/d7030295

Fox J, Weisberg S (2019) An R Companion to Applied Regression, Third. Sage, Thousand Oaks (CA)

Frankham R (2012) How closely does genetic diversity in finite populations conform to predictions of neutral theory? Large deficits in regions of low recombination. Heredity 108:167–178. 10.1038/hdy.2011.66

Frankham R (1995) Effective population size/adult population size ratios in wildlife: a review. Genet Res 66:95–107. 10.1017/S0016672300034455

Fuentes-Pardo AP, Ruzzante DE (2017) Whole-genome sequencing approaches for conservation biology: Advantages, limitations and practical recommendations. Molecular Ecology 26:5369–5406. 10.1111/mec.14264

Gargiulo R, Budde KB, Heuertz M (2024) Mind the lag: understanding genetic extinction debt for conservation. Trends in Ecology & Evolution 0: 10.1016/j.tree.2024.10.008

Garner BA, Hoban S, Luikart G (2020) IUCN Red List and the value of integrating genetics. Conserv Genet 21:795–801. 10.1007/s10592-020-01301-6

Gopalakrishnan S, Samaniego Castruita JA, Sinding M-HS, et al (2017) The wolf reference genome sequence (Canis lupus lupus) and its implications for Canis spp. population genomics. BMC Genomics 18:495. 10.1186/s12864-017-3883-3

Grundler MR, Singhal S, Cowan MA, Rabosky DL (2019) Is genomic diversity a useful proxy for census population size? Evidence from a species-rich community of desert lizards. Molecular Ecology 28:1664–1674. 10.1111/mec.15042

Hague MTJ, Routman EJ (2016) Does population size affect genetic diversity? A test with sympatric lizard species. Heredity 116:92–98. 10.1038/hdy.2015.76

Hoban S, Bruford M, D’Urban Jackson J, et al (2020) Genetic diversity targets and indicators in the CBD post-2020 Global Biodiversity Framework must be improved. Biological Conservation 248:108654. 10.1016/j.biocon.2020.108654

Hoban S, da Silva JM, Hughes A, et al (2024) Too simple, too complex, or just right? Advantages, challenges, and guidance for indicators of genetic diversity. BioScience 74:269–280. 10.1093/biosci/biae006

Huffmeyer AA, Sikich JA, Vickers TW, et al (2022) First reproductive signs of inbreeding depression in Southern California male mountain lions (*Puma concolor*). Theriogenology 177:157–164. 10.1016/j.theriogenology.2021.10.016

IUCN (2023) The IUCN Red List of Threatened Species Version 23-1. In: IUCN Red List of Threatened Species. https://www.iucnredlist.org/en. Accessed 27 Jan 2024

IUCN Standards and Petitions Committee (2024) Guidelines for Using the IUCN Red List Categories and Criteria: Version 16

Jeon JY, Black A, Heenkenda E, et al (2024) Genomic Diversity as a Key Conservation Criterion: Proof- of-Concept From Mammalian Whole-Genome Resequencing Data. Evolutionary Applications 17:. 10.1111/eva.70000

Jhala YV, Banerjee K, Chakrabarti S, et al (2019) Asiatic Lion: Ecology, Economics, and Politics of Conservation. Front Ecol Evol 7:. 10.3389/fevo.2019.00312

Johnson WE, Onorato DP, Roelke ME, et al (2010) Genetic Restoration of the Florida Panther. Science 329:1641–1645. 10.1126/science.1192891

Jorge MLSP, Galetti M, Ribeiro MC, Ferraz KMPMB (2013) Mammal defaunation as surrogate of trophic cascades in a biodiversity hotspot. Biological Conservation 163:49–57. 10.1016/j.biocon.2013.04.018

José R. Castelló (2020) Felids and Hyenas of the World. Princeton University Press, New Jersey

Kassambara A (2023) ggpubr: “ggplot2” Based Publication Ready Plots

Kim S, Cho YS, Kim H-M, et al (2016) Comparison of carnivore, omnivore, and herbivore mammalian genomes with a new leopard assembly. Genome Biology 17:211. 10.1186/s13059-016-1071-4

Kitchener AC, Breitenmoser-Würsten C, Eizirik E, et al (2017) A revised taxonomy of the Felidae : The final report of the Cat Classification Task Force of the IUCN Cat Specialist Group

Kuderna LFK, Gao H, Janiak MC, et al (2023) A global catalog of whole-genome diversity from 233 primate species. Science 380:906–913. 10.1126/science.abn7829

Laikre L, Hoban S, Bruford MW, et al (2020) Post-2020 goals overlook genetic diversity. Science 367:1083–1085. 10.1126/science.abb2748

Leffler EM, Bullaughey K, Matute DR, et al (2012) Revisiting an Old Riddle: What Determines Genetic Diversity Levels within Species? PLOS Biology 10:e1001388. 10.1371/journal.pbio.1001388

Lescroart J, Bonilla-Sánchez A, Napolitano C, et al (2023) Extensive Phylogenomic Discordance and the Complex Evolutionary History of the Neotropical Cat Genus Leopardus. Molecular Biology and Evolution 40:msad255. 10.1093/molbev/msad255

Li H, Durbin R (2009) Fast and accurate short read alignment with Burrows–Wheeler transform. Bioinformatics 25:1754–1760. 10.1093/bioinformatics/btp324

Lin M, Escalona M, Sahasrabudhe R, et al (2022) A Reference Genome Assembly of the Bobcat, Lynx rufus. Journal of Heredity esac031. 10.1093/jhered/esac031

Lino A, Fonseca C, Rojas D, et al (2019) A meta-analysis of the effects of habitat loss and fragmentation on genetic diversity in mammals. Mammalian Biology 94:69–76. 10.1016/j.mambio.2018.09.006

Liu Y-C, Sun X, Driscoll C, et al (2018) Genome-Wide Evolutionary Analysis of Natural History and Adaptation in the World’s Tigers. Current Biology 28:3840–3849.e6. 10.1016/j.cub.2018.09.019

Lorenzana GP, Figueiró HV, Kaelin CB, et al (2022) Whole-genome sequences shed light on the demographic history and contemporary genetic erosion of free-ranging jaguar (Panthera onca) populations. Journal of Genetics and Genomics 49:77–80. 10.1016/j.jgg.2021.10.006

Lyons SK, Smith FA, Ernest SKM (2019) Macroecological patterns of mammals across taxonomic, spatial, and temporal scales. Journal of Mammalogy 100:1087–1104. 10.1093/jmammal/gyy171

Macdonald EA, Burnham D, Hinks AE, et al (2015) Conservation inequality and the charismatic cat: Felis felicis. Global Ecology and Conservation 3:851–866. 10.1016/j.gecco.2015.04.006

Mackintosh A, Laetsch DR, Hayward A, et al (2019) The determinants of genetic diversity in butterflies. Nat Commun 10:3466. 10.1038/s41467-019-11308-4

Markert JA, Champlin DM, Gutjahr-Gobell R, et al (2010) Population genetic diversity and fitness in multiple environments. BMC Evolutionary Biology 10:205. 10.1186/1471-2148-10-205

Martin AP, Palumbi SR (1993) Body size, metabolic rate, generation time, and the molecular clock. Proc Natl Acad Sci USA 90:4087–4091. 10.1073/pnas.90.9.4087

Myhre AM, Engen S, Sæther B-E (2016) Effective size of density-dependent populations in fluctuating environments. Evolution 70:2431–2446. 10.1111/evo.13063

Nabholz B, Mauffrey J-F, Bazin E, et al (2008) Determination of Mitochondrial Genetic Diversity in Mammals. Genetics 178:351–361. 10.1534/genetics.107.073346

Nielsen C, Thompson D, Kelly M, Lopez-Gonzalez CA (2016) Puma concolor. The IUCN Red List of Threatened Species 2015 2016:. 10.2305/IUCN.UK.2015-4.RLTS.T18868A50663436.en

O’Brien SJ, Johnson WE, Driscoll CA, et al (2017) Conservation Genetics of the Cheetah: Lessons Learned and New Opportunities. Journal of Heredity 108:671–677. 10.1093/jhered/esx047

Otto SP (2018) Adaptation, speciation and extinction in the Anthropocene. Proceedings of the Royal Society B: Biological Sciences 285:20182047. 10.1098/rspb.2018.2047

Paijmans JLA, Barlow A, Becker MS, et al (2021) African and Asian leopards are highly differentiated at the genomic level. Current Biology 31:1872–1882.e5. 10.1016/j.cub.2021.03.084

Palstra FP, Ruzzante DE (2008) Genetic estimates of contemporary effective population size: what can they tell us about the importance of genetic stochasticity for wild population persistence? Molecular Ecology 17:3428–3447. 10.1111/j.1365-294X.2008.03842.x

Pečnerová P, Garcia-Erill G, Liu X, et al (2021) High genetic diversity and low differentiation reflect the ecological versatility of the African leopard. Current Biology 31:1862–1871.e5. 10.1016/j.cub.2021.01.064

Pérez-Irineo G, Santos-Moreno A (2013) TRENDS IN RESEARCH ON TERRESTRIAL SPECIES OF THE ORDER CARNIVORA

Perry GH, Melsted P, Marioni JC, et al (2012) Comparative RNA sequencing reveals substantial genetic variation in endangered primates. Genome Res 22:602–610. 10.1101/gr.130468.111

Petit RA, Hall MB, Tonkin-Hill G, et al (2023) fastq-dl: efficiently download FASTQ files from SRA or ENA repositories

Pimm SL, Dollar L, Bass Jr OL (2006) The genetic rescue of the Florida panther. Animal Conservation 9:115–122. 10.1111/j.1469-1795.2005.00010.x

Prasad A, Lorenzen ED, Westbury MV (2022) Evaluating the role of reference-genome phylogenetic distance on evolutionary inference. Molecular Ecology Resources 22:45–55. 10.1111/1755-0998.13457

Prentice MB, Bowman J, Murray DL, et al (2019) Evaluating evolutionary history and adaptive differentiation to identify conservation units of Canada lynx (*Lynx canadensis*). Global Ecology and Conservation 20:e00708. 10.1016/j.gecco.2019.e00708

Prost S, Machado AP, Zumbroich J, et al (2022) Genomic analyses show extremely perilous conservation status of African and Asiatic cheetahs (Acinonyx jubatus). Molecular Ecology 31:4208–4223. 10.1111/mec.16577

Quinlan AR, Hall IM (2010) BEDTools: a flexible suite of utilities for comparing genomic features. Bioinformatics 26:841–842. 10.1093/bioinformatics/btq033

Ramirez JL, Lescroart J, Figueiró HV, et al (2022) Genomic Signatures of Divergent Ecological Strategies in a Recent Radiation of Neotropical Wild Cats. Molecular Biology and Evolution 39:msac117. 10.1093/molbev/msac117

Reed DH, Frankham R (2003) Correlation between Fitness and Genetic Diversity. Conservation Biology 17:230–237. 10.1046/j.1523-1739.2003.01236.x

Riley SPD, Serieys LEK, Pollinger JP, et al (2014) Individual Behaviors Dominate the Dynamics of an Urban Mountain Lion Population Isolated by Roads. Current Biology 24:1989–1994. 10.1016/j.cub.2014.07.029

Ripple WJ, Beschta RL (2006) Linking a cougar decline, trophic cascade, and catastrophic regime shift in Zion National Park. Biological Conservation 133:397–408. 10.1016/j.biocon.2006.07.002

Ripple WJ, Beschta RL (2008) Trophic cascades involving cougar, mule deer, and black oaks in Yosemite National Park. Biological Conservation 141:1249–1256. 10.1016/j.biocon.2008.02.028

Romiguier J, Gayral P, Ballenghien M, et al (2014) Comparative population genomics in animals uncovers the determinants of genetic diversity. Nature 515:261–263. 10.1038/nature13685

Ros-Freixedes R, Battagin M, Johnsson M, et al (2018) Impact of index hopping and bias towards the reference allele on accuracy of genotype calls from low-coverage sequencing. Genetics Selection Evolution 50:64. 10.1186/s12711-018-0436-4

Sarasola JH, Zanón-Martínez JI, Costán AS, Ripple WJ (2016) Hypercarnivorous apex predator could provide ecosystem services by dispersing seeds. Sci Rep 6:19647. 10.1038/srep19647

Saremi NF, Supple MA, Byrne A, et al (2019) Puma genomes from North and South America provide insights into the genomic consequences of inbreeding. Nat Commun 10:4769. 10.1038/s41467-019-12741-1

Schloerke B, Cook D, Larmaran J, et al (2024) GGally: Extension to “ggplot2”

Schmidt AF, Finan C (2018) Linear regression and the normality assumption. Journal of Clinical Epidemiology 98:146–151. 10.1016/j.jclinepi.2017.12.006

Schmidt C, Hoban S, Hunter M, et al (2023) Genetic diversity and IUCN Red List status. Conservation Biology 37:e14064. 10.1111/cobi.14064

Schwartz MK, Mills LS, Ortega Y, et al (2003) Landscape location affects genetic variation of Canada lynx (Lynx canadensis). Molecular Ecology 12:1807–1816. 10.1046/j.1365-294X.2003.01878.x

Silva M, Brimacombe M, Downing JA (2001) Effects of body mass, climate, geography, and census area on population density of terrestrial mammals. Global Ecology and Biogeography 10:469–485. 10.1046/j.1466-822x.2001.00261.x

Slate J, Kruuk LEB, Marshall TC, et al (2000) Inbreeding depression influences lifetime breeding success in a wild population of red deer (Cervus elaphus). Proceedings of the Royal Society of London Series B: Biological Sciences 267:1657–1662. 10.1098/rspb.2000.1192

Smith FA, Lyons SK (2013) Macroecological Patterns of Body Size in Mammals across Time and Space. In: Animal Body Size: Linking Pattern and Process across Space, Time, and Taxonomic Group. University of Chicago Press, pp 116–144

Solari KA, Morgan S, Poyarkov AD, et al (2023) Extreme in Every Way: Exceedingly Low Genetic Diversity in Snow Leopards Due to Persistently Small Population Size. 2023.12.14.571340

Sousa V, Hey J (2013) Understanding the origin of species with genome-scale data: modelling gene flow. Nat Rev Genet 14:404–414. 10.1038/nrg3446

Takahashi Y, Tanaka R, Yamamoto D, et al (2018) Balanced genetic diversity improves population fitness. Proceedings of the Royal Society B: Biological Sciences 285:20172045. 10.1098/rspb.2017.2045

Teixeira JC, Huber CD (2021) The inflated significance of neutral genetic diversity in conservation genetics. Proceedings of the National Academy of Sciences 118:e2015096118. 10.1073/pnas.2015096118

Tensen L (2018a) Biases in wildlife and conservation research, using felids and canids as a case study. Global Ecology and Conservation 15:e00423. 10.1016/j.gecco.2018.e00423

Tensen L (2018b) Species characteristics of felids and canids, and the number of articles published for each species between 2013 and 2017. Data in Brief 21:201–211. 10.1016/j.dib.2018.09.132

Theissinger K, Fernandes C, Formenti G, et al (2023) How genomics can help biodiversity conservation. Trends in Genetics 39:545–559. 10.1016/j.tig.2023.01.005

Väli Ü, Einarsson A, Waits L, Ellegren H (2008) To what extent do microsatellite markers reflect genome-wide genetic diversity in natural populations? Molecular Ecology 17:3808–3817. 10.1111/j.1365-294X.2008.03876.x

White EP, Ernest SKM, Kerkhoff AJ, Enquist BJ (2007) Relationships between body size and abundance in ecology. Trends in Ecology & Evolution 22:323–330. 10.1016/j.tree.2007.03.007

Wickham H (2016) ggplot2: Elegant Graphics for Data Analyisis. Springer-Verlag, New York

Wilder AP, Supple MA, Subramanian A, et al (2023) The contribution of historical processes to contemporary extinction risk in placental mammals. Science 380:eabn5856. 10.1126/science.abn5856

Willi Y, Kristensen TN, Sgrò CM, et al (2022) Conservation genetics as a management tool: The five best-supported paradigms to assist the management of threatened species. Proceedings of the National Academy of Sciences 119:e2105076119. 10.1073/pnas.2105076119

Willig M, Lyons S, Stevens R (2009) Spatial methods for the macroecological study of bats

Willoughby JR, Sundaram M, Wijayawardena BK, et al (2015) The reduction of genetic diversity in threatened vertebrates and new recommendations regarding IUCN conservation rankings. Biological Conservation 191:495–503. 10.1016/j.biocon.2015.07.025

Wilson DE, Mittermeier RA (2009) Handbook of the Mammals of the World Volume 1: Carnivores, 1st edn. Lynx Edicions, Barcelona

Wooten MC, Smith MH (1985) LARGE MAMMALS ARE GENETICALLY LESS VARIABLE? Evolution 39:210–212. 10.1111/j.1558-5646.1985.tb04095.x

Yuan J, Kitchener AC, Lackey LB, et al (2024) The genome of the black-footed cat: Revealing a rich natural history and urgent conservation priorities for small felids. Proceedings of the National Academy of Sciences 121:e2310763120. 10.1073/pnas.2310763120

Yuan J, Wang G, Zhao L, et al (2023) How genomic insights into the evolutionary history of clouded leopards inform their conservation. Science Advances 9:eadh9143. 10.1126/sciadv.adh9143

Zanin M, Palomares F, Brito D (2015) What we (don’t) know about the effects of habitat loss and fragmentation on felids. Oryx 49:96–106. 10.1017/S0030605313001609

