## Supplemental Figures for "Genomic diversity in felids correlates with range and density, not census size"

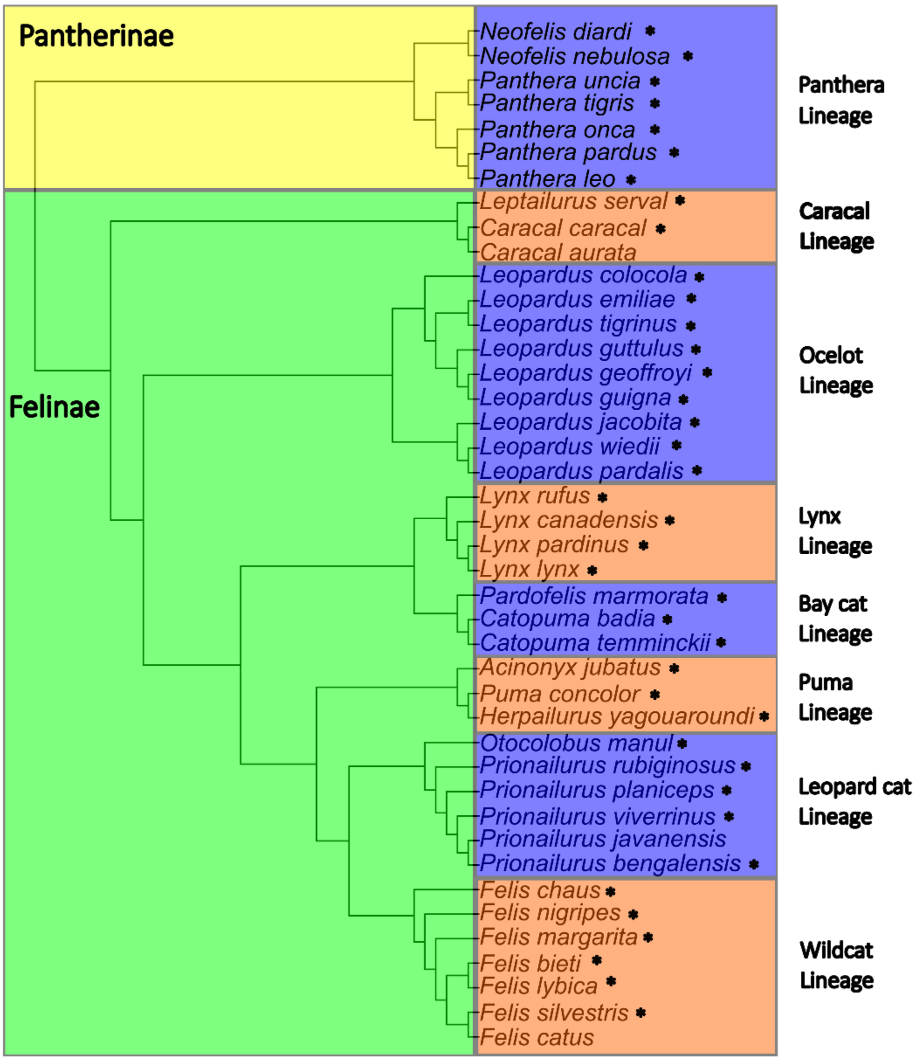


**Fig. S1** Phylogenetic tree of the Felidae family. Reproduced from the nuclear phylogeny in Li et al. (2016), with addition of *Leopardus emiliae*. Branch length of the split between *Leopardus emiliae* and *L. tigrinus* was arbitrarily set at 1. Species included in our dataset are marked with an asterisk

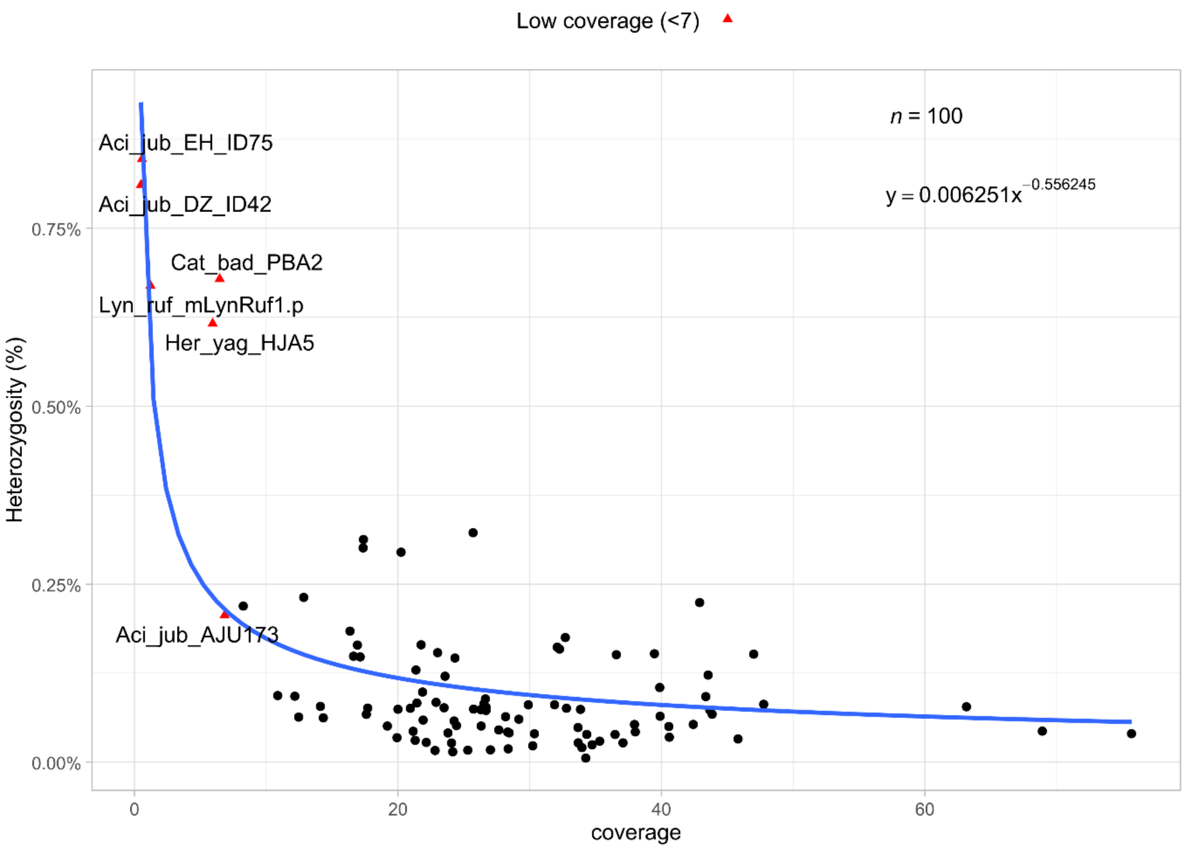


**Fig. S2** Power regression for the relationship between estimates of heterozygosity and sample coverage. Red triangles denote datapoints with low coverage (< 7) and were removed from the dataset for the subsequent analyses

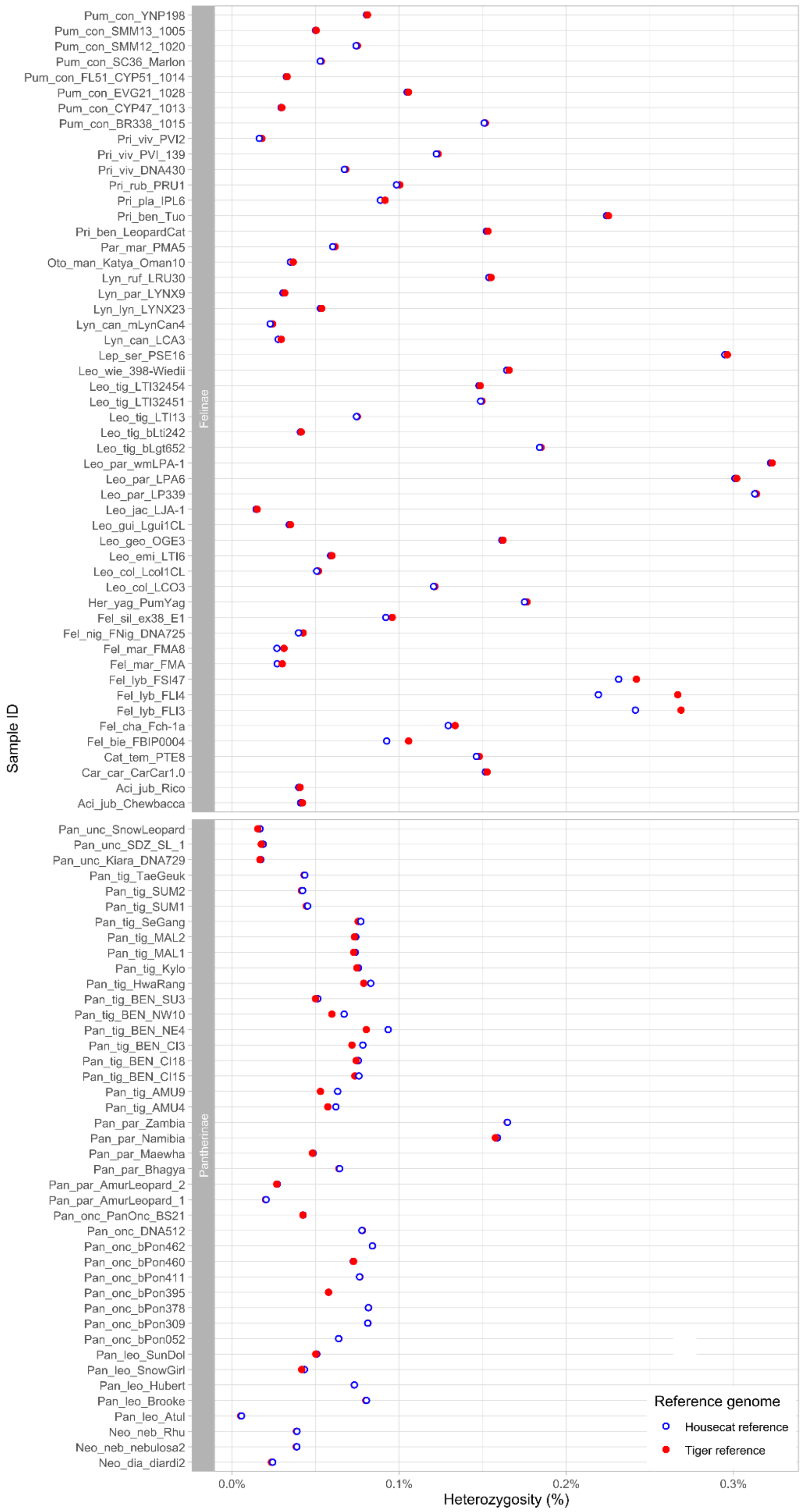


**Fig. S3** Per sample heterozygosity estimates for two reference genomes. Samples are grouped per subfamily. Blue open circles denote estimates made with the housecat reference; red filled circles denote estimates made with the tiger reference

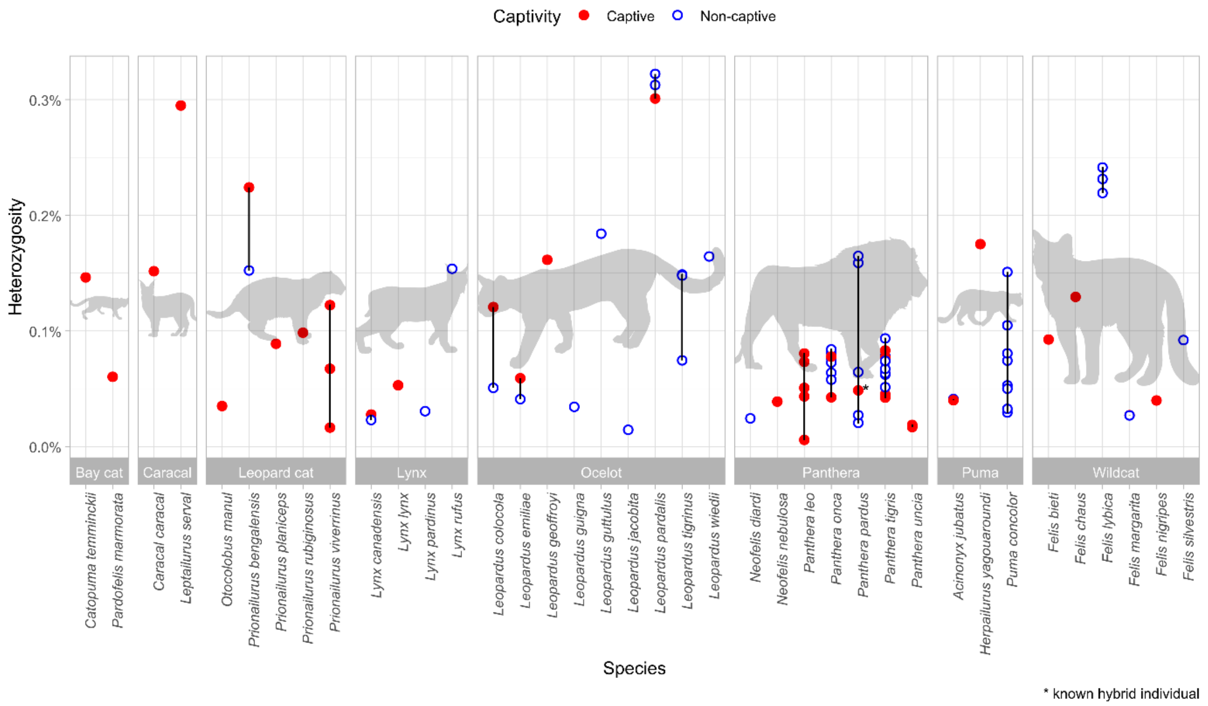


**Fig. S4** Estimates of heterozygosity for all samples using the cat reference genome, grouped per lineage and species. Red filled circles denote samples from captive populations; blue open circles denote samples from wild populations. Silhouettes obtained from PhyloPic (www.phylopic.org)

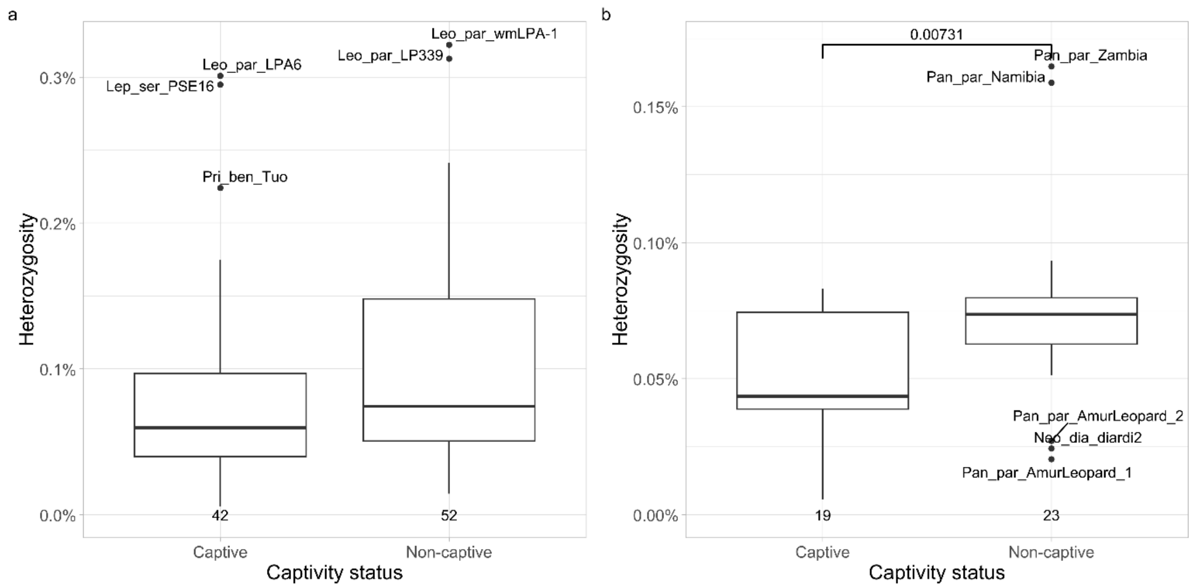


**Fig. S5** Comparison of heterozygosity between captive and non-captive samples **a** across the entire dataset, **b** within the Panthera lineage. Significance tests were performed using a Welch's t-test

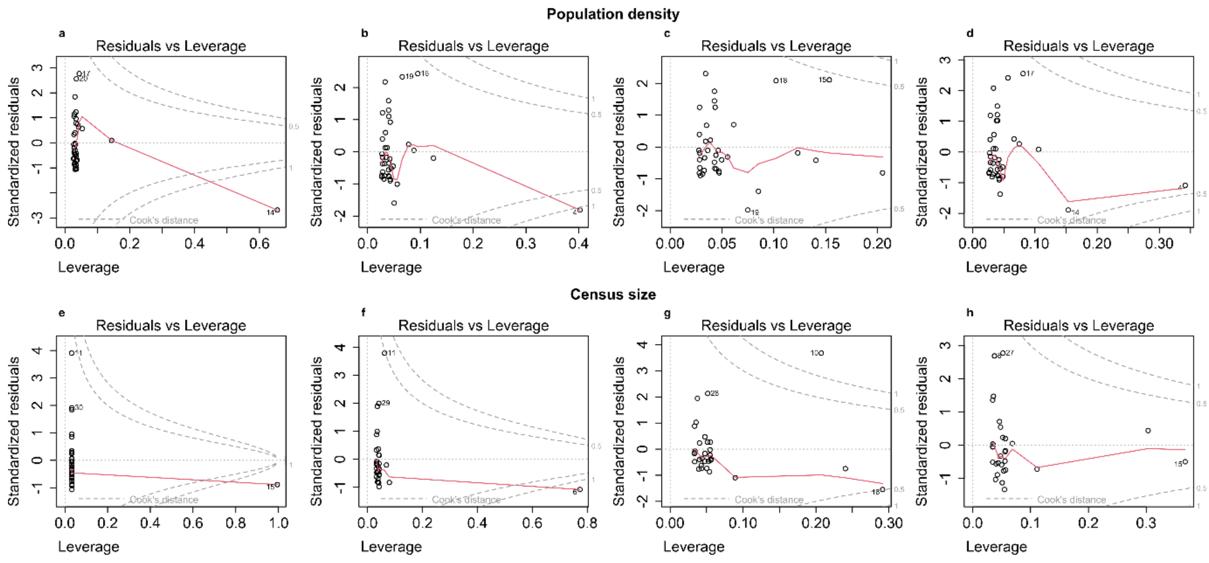


**Fig. S6** Cook's distance plots to identify outliers in the mean population density (**a-d**) and census size (**e-h**) data at subspecies level. Cook's distance plot for **a** unfiltered population density data, **b** population density data excluding *Leopardus guigna,* **c** population density data excluding *Felis chaus,* **d** population density data with a more conservative value for *L. guigna,* **e** unfiltered census size data, **f** census size data excluding *Lynx rufus,* **g** census size data excluding *Felis silvestris,* **h** census size data excluding *Leopardus pardalis*

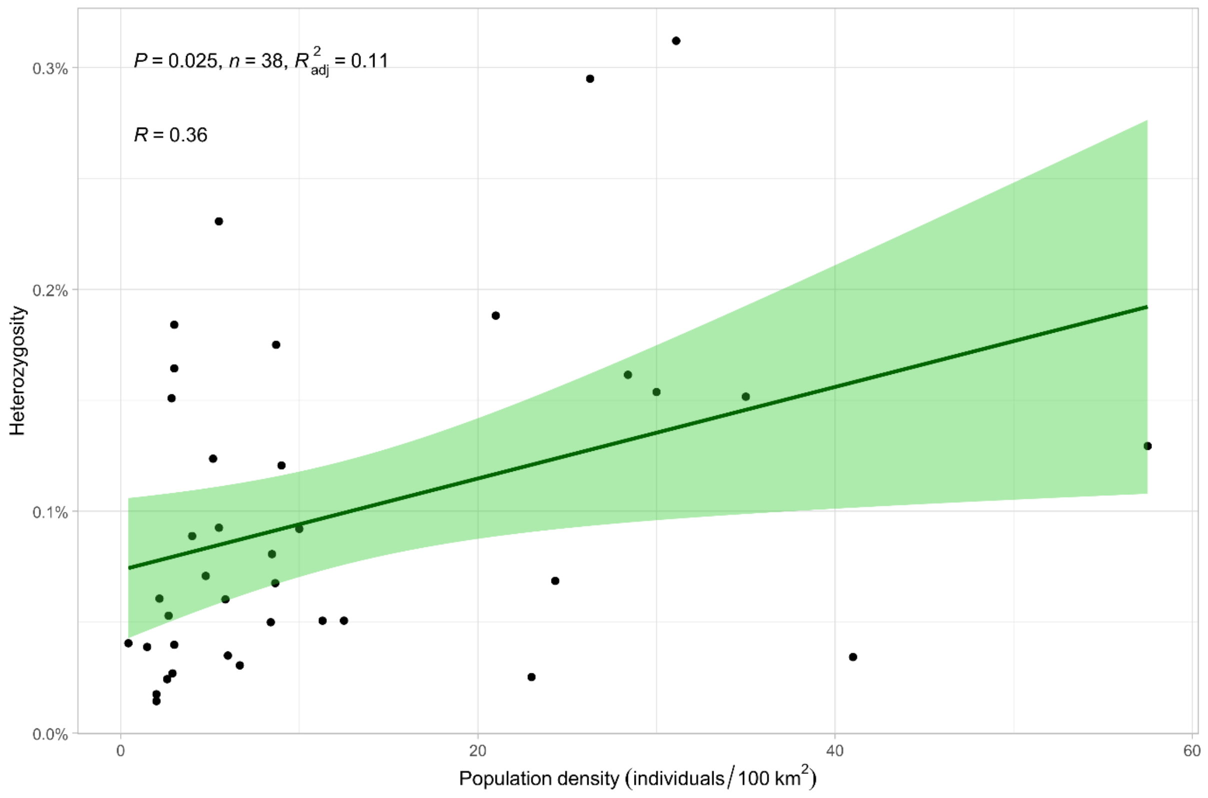


**Fig. S7** Linear regression between heterozygosity and population density using a more conservative value for *Leopardus guigna*

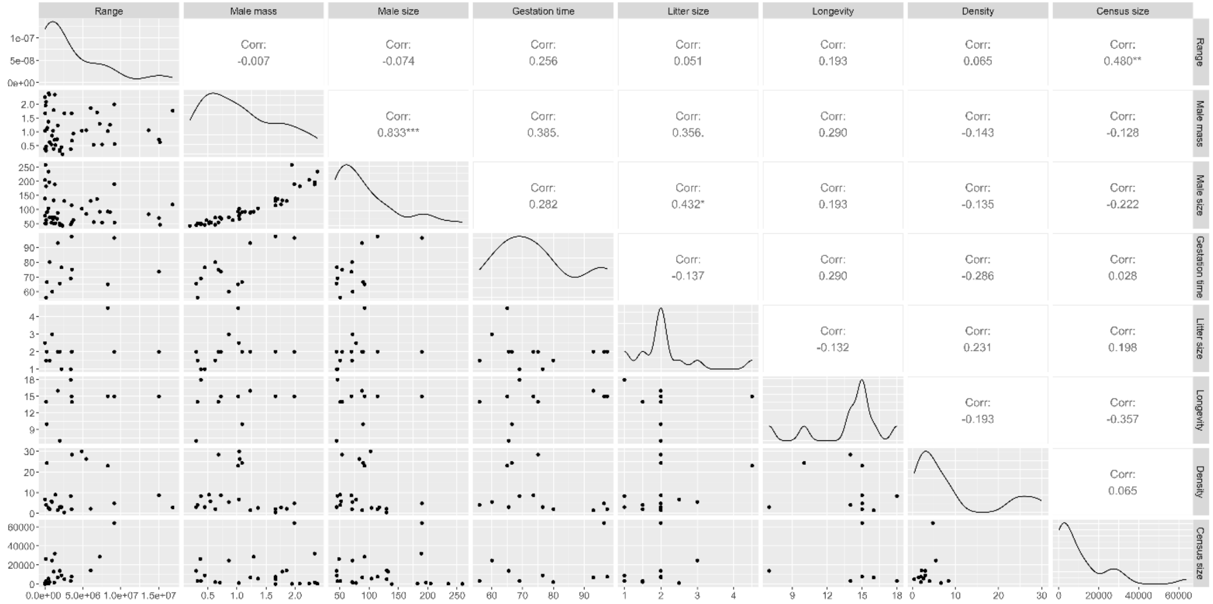

**Fig. S8** Correlation matrix of independent variables at the subspecies level using Kendall's τ

### References

Li G, Davis BW, Eizirik E, Murphy WJ (2016) Phylogenomic evidence for ancient hybridization in the genomes of living cats (Felidae). Genome Res 26:1–11. https://doi.org/10.1101/gr.186668.114
